# Structure of alpha-synuclein fibrils derived from human Lewy body dementia tissue

**DOI:** 10.1101/2023.01.09.523303

**Authors:** Dhruva D. Dhavale, Alexander M. Barclay, Collin G. Borcik, Katherine Basore, Isabelle R. Gordon, Jialu Liu, Moses H. Milchberg, Jennifer O’shea, Michael J. Rau, Zachary Smith, Soumyo Sen, Brock Summers, John Smith, Owen A. Warmuth, Qian Chen, James A. J. Fitzpatrick, Charles D. Schwieters, Emad Tajkhorshid, Chad M. Rienstra, Paul T. Kotzbauer

## Abstract

The defining feature of Parkinson disease (PD) and Lewy body dementia (LBD) is the accumulation of alpha-synuclein (Asyn) fibrils in Lewy bodies and Lewy neurites. We developed and validated a novel method to amplify Asyn fibrils extracted from LBD postmortem tissue samples and used solid state nuclear magnetic resonance (SSNMR) studies to determine atomic resolution structure. Amplified LBD Asyn fibrils comprise two protofilaments with pseudo-2_1_ helical screw symmetry, very low twist and an interface formed by antiparallel beta strands of residues 85-93. The fold is highly similar to the fold determined by a recent cryo-electron microscopy study for a minority population of twisted single protofilament fibrils extracted from LBD tissue. These results expand the structural landscape of LBD Asyn fibrils and inform further studies of disease mechanisms, imaging agents and therapeutics targeting Asyn.

## Main

Parkinson disease (PD) is diagnosed clinically based on motor signs that include tremor, bradykinesia, rigidity and impaired postural reflexes. PD is defined pathologically by the accumulation of alpha-synuclein (Asyn) fibrils in neuronal cytoplasmic and neuritic inclusions known as Lewy bodies (LBs) and Lewy neurites (LNs)^1^. The role of Asyn in the pathogenesis of PD is supported by the identification of dominant mutations in the gene encoding Asyn (*SNCA*) in rare familial versions of PD^2-9^. Dementia occurs frequently in PD and is associated with pathologic Asyn accumulation in neocortex^10-12^. Dementia sometimes begins at approximately the same time as motor symptoms (Dementia with Lewy bodies or DLB), or up to 20 years after motor symptoms begin (PD with dementia or PDD). Lewy body dementia (LBD) encompasses this spectrum of clinical presentations.

Analysis of Asyn fibril structure in LBD guides the identification of high affinity ligands needed to develop a Positron Emission Tomography (PET) imaging agent that can quantify the accumulation of Asyn fibrils in living individuals. An Asyn imaging agent would improve diagnosis and provide a biomarker for tracking disease progression. It would also enable assessment of target engagement for therapeutic strategies targeting Asyn^13^. Structural analysis of Asyn fibrils may also provide insight into disease mechanisms, including (1) promotion of nucleation, growth and clearance of fibrils; and (2) regulation of cell-to-cell transfer that underlies disease progression.

Different Asyn fibril polymorphs are produced by incubating human recombinant monomeric Asyn protein under different conditions to induce fibril formation, including differences in pH, buffers and salts, as demonstrated by SSNMR^14-21^ and cryo-EM studies^22-36^. The structure of Asyn fibrils isolated from postmortem multiple system atrophy (MSA) brain tissue, where fibrils accumulate in oligodendroglial cells, has been determined by cryo-EM^37^. The fold of MSA Asyn fibrils is distinct from in vitro forms, although it does share one structural element, a Greek key fold, with a subset of the reported in vitro forms^16,37^. LBD Asyn fibril structure has been difficult to analyze with cryo-EM because of the very low twist present in fibrils isolated from tissue. Yang et al recently reported a 2.2 Å structure (PDB 8A9L) of Asyn fibrils from a minority population (25%) of LBD fibrils found to have high twist and to be amenable to cryo-EM analysis after extraction from LBD tissue^38^. The fold of these single protofilament LBD fibrils, termed the ‘Lewy fold’, is distinct from structures previously reported for MSA and for Asyn fibrils assembled in vitro.

To analyze LBD fibril structure by SSNMR, we developed a novel method to amplify fibrils extracted from postmortem LBD brain tissue using isotopically labeled human recombinant Asyn. We utilized seeding properties in a cell culture system to guide method development by comparing the properties of amplified fibrils with tissue-derived fibrils. The amplification method enabled us to prepare multiple milligram quantities of amplified fibrils, which were used for structure determination by SSNMR. LBD amplified fibrils comprise two protofilaments with pseudo-2_1_ helical screw symmetry, very low twist and an interface formed by antiparallel beta strands of residues 85-93. The fold of the LBD amplified protofilaments is highly similar to the fold of the single protofilament fibrils with high twist analyzed by single-particle cryo-EM.

## Results

### Production of amplified fibrils from LBD and MSA postmortem tissue

Previous studies of fibril growth for WT and mutant Asyn fibrils indicate that fibril structure is templated during growth of fibril seeds in the presence of recombinant Asyn monomer^39^. To produce fibrils that replicate the structure present in LBD tissue, we developed a protocol to isolate fibrils from brain tissue samples in an insoluble fraction and use them as seeds to grow fibrils with recombinant human Asyn protein produced in E. Coli. This amplification protocol utilizes multiple cycles of sonication-mediated fragmentation followed by quiescent incubation for 2-3 days (Fig. 1a).

**Fig. 1:**
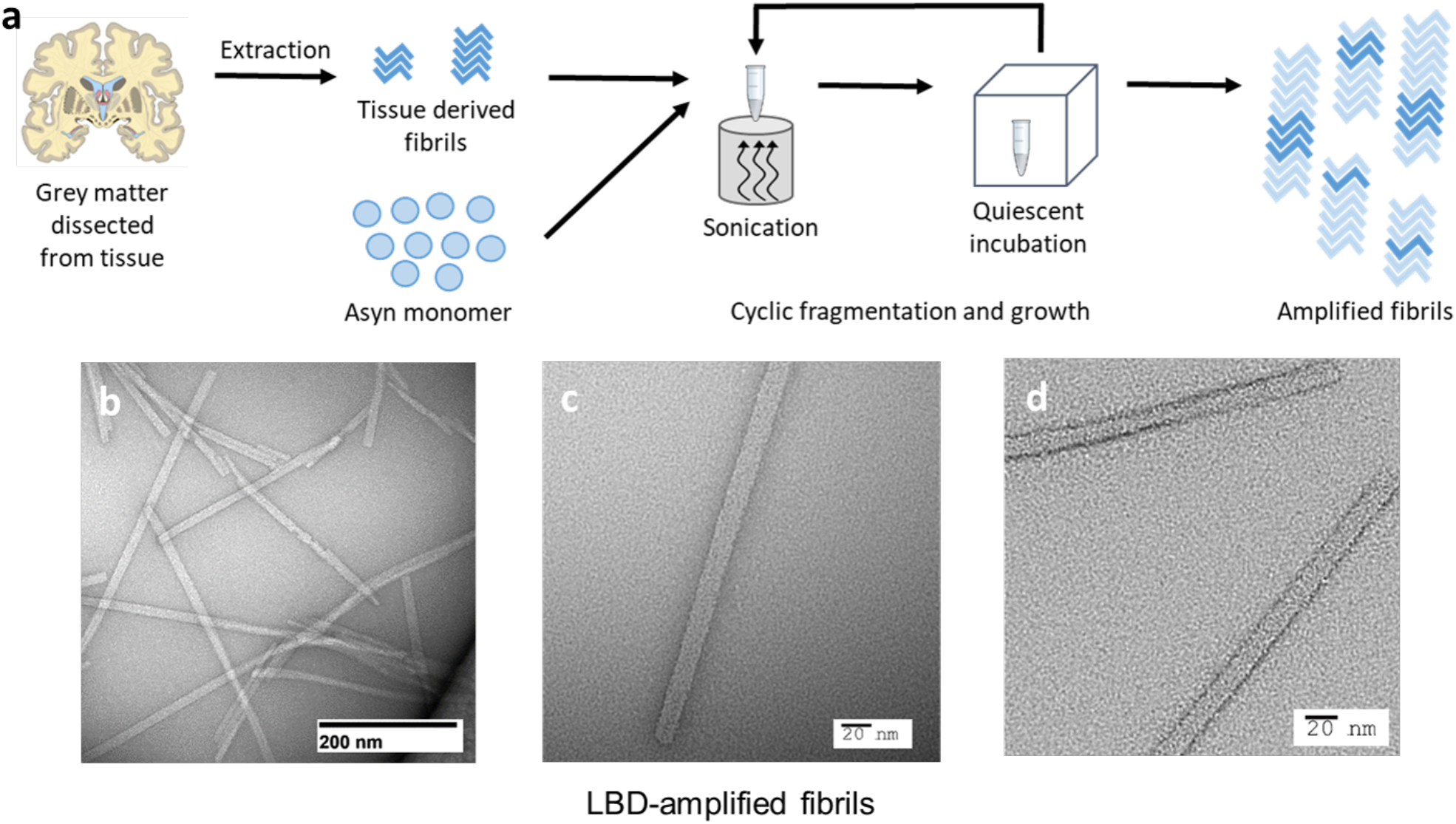
Schematic illustrating extraction and amplification of LBD Asyn fibrils derived from postmortem tissue samples. **a**, Fibrils were isolated in an insoluble fraction by sequential extraction and centrifugation of tissue samples. The insoluble fraction seeds were combined with recombinant Asyn monomer and subjected to multiple rounds of sonication followed by quiescent incubation at 37 °C to generate amplified fibrils. **b-d**, Representative negative stain TEM images of LBD amplified fibrils.

We used a highly sensitive radioligand binding assay to quantify LBD fibril amplification and guide the optimization of the protocol. We used the binding site density (B_max_ = 42 pmol/nmol) and affinity (K_d_ = 5 nM) of the radioligand for LBD amplified fibrils to convert measurements of bound radioligand in binding assays to fibril concentration (µM)^40,41^. An important goal was to optimize amplification from fibril seeds isolated from LBD tissue samples, while minimizing the accumulation of fibrils with insoluble fractions prepared from control tissue samples, which were selected based on the absence of pathologic Asyn as determined by immunohistochemistry.

LBD amplification was initially optimized with natural abundance Asyn monomer. At the end of 6 cycles, we observed 16-fold to 35-fold greater mass of fibrils amplified from LBD samples compared to control samples (Supplementary Table 1). We further optimized amplification in the presence of isotopically labeled Asyn monomer to facilitate SSNMR studies and selected case LBD1 for large scale amplifications. We observed 11-fold and 18-fold greater mass of LBD1 fibrils compared to two control cases (Supplementary Tables 2 and 3). We used a micro-BCA protein concentration assay to further evaluate fibril concentration and confirm the specificity of amplification for LBD compared to control samples at the end of the 6^th^ cycle (Supplementary Table 4). We obtained a yield of 5 mg and 3.8 mg of LBD1 fibrils amplified with uniform [^13^C, ^15^N] labeled Asyn monomer (uCN) and with uniform [^2^H, ^13^C, ^15^N] labeled Asyn monomer (uDCN) respectively, which were utilized for SSNMR studies (Supplementary Table 4).

Negative-stain transmission EM analysis of amplified LBD fibrils showed straight fibrils with a diameter of 10-18 nm and no visible twist (Fig. 1b,d). Fibrils amplified from control tissue samples had smaller diameter and more curvature (Extended Data Fig. 1a,b). As an approach to further evaluate structural fidelity of amplification, we also amplified fibrils from MSA tissue samples, which appear twisted (Extended Data Fig. 1c). We also compared the morphology of amplified fibrils to those assembled in vitro by incubation of recombinant Asyn monomer at 37 °C with continuous shaking^39,41-47^. These fibrils appear distinct from all forms of amplified fibrils, with paired helical filament morphology (Extended Data Fig. 1d).

### Characterization of amplified fibrils by seeding Asyn aggregation in HEK293 cells – support for the structural fidelity of fibril amplification

We previously used a biosensor cell line to evaluate seeding activity of fibrils extracted from postmortem brain tissue samples and observed distinct seeding properties for LBD and MSA tissue^47^. We seeded these biosensor cells with fibrils amplified from either LBD or MSA tissue samples and compared their seeding properties to fibrils extracted directly from LBD and MSA tissue. We observed a consistent morphologic difference for intracellular inclusions produced by seeding with LBD and MSA amplified fibrils (after 4^th^ cycle) in HEK293 cells (Fig. 2a-d). Inverted fluorescence microscopy images show that inclusions seeded with LBD amplified fibrils consistently have a more compact, clumped appearance. In contrast, inclusions seeded with MSA amplified fibrils contain long filamentous strands that expand throughout the cytoplasm. This distinct morphological appearance recapitulates the morphology produced by seeding with insoluble fractions obtained from LBD and MSA tissue samples (Fig. 2)^47^. Similar morphologic features were observed after 4 and 6 cycles of amplification (data not shown), and the morphology was consistent between multiple cases of the same disease phenotype.

**Fig. 2:**
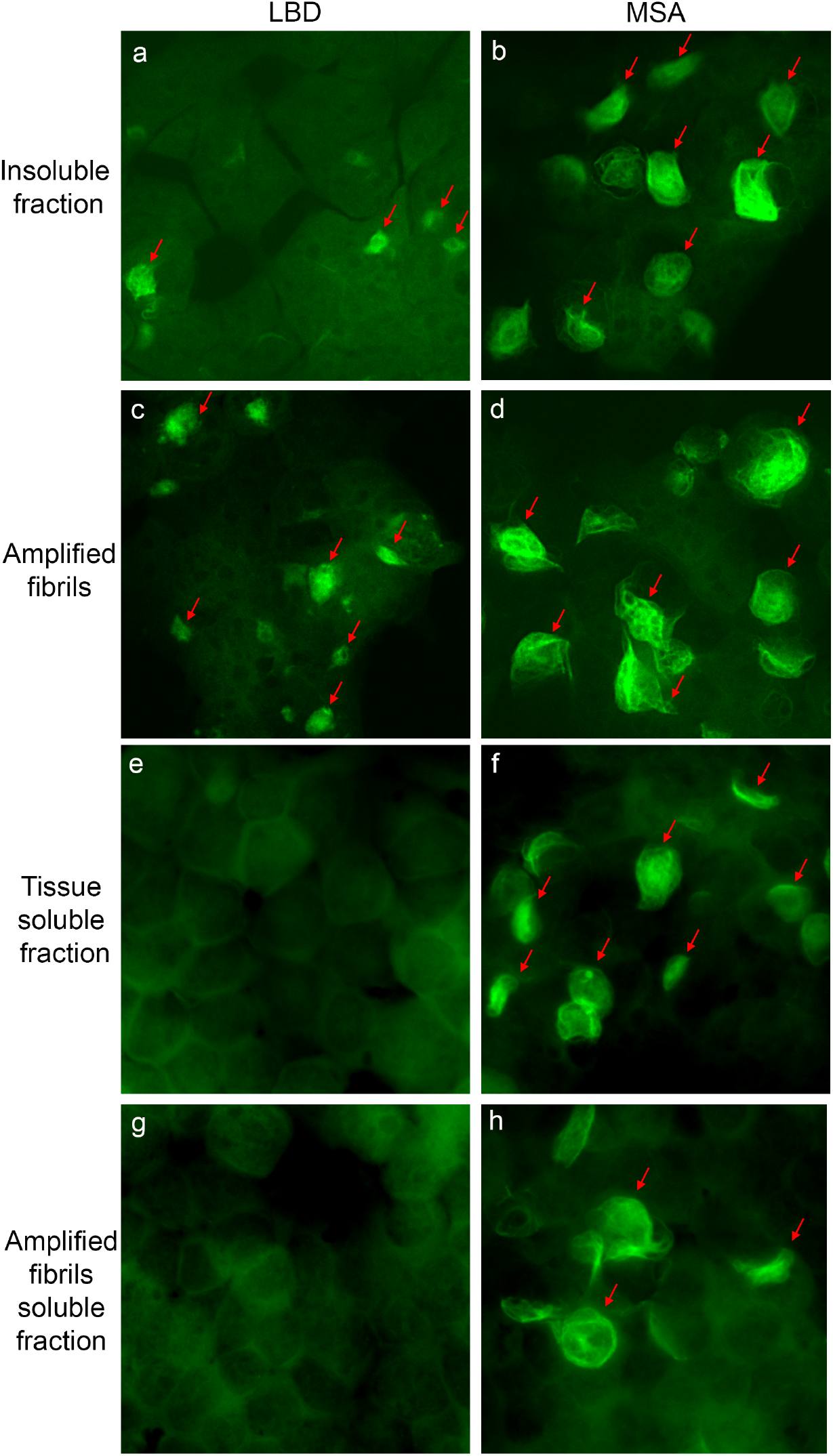
Seeding properties of tissue-derived and amplified fibrils in HEK 293 biosensor cells expressing A53T Asyn-CFP/YFP. Fluorescence images show cells seed with: **a**, LBD-Insoluble fraction. **b**, MSA-insoluble fraction. **c**, LBD-amplified fibrils. **d**, MSA-amplified fibrils. **e**, LBD-tissue soluble fraction. **f**, MSA-tissue soluble fraction. **g**, LBD-amplified fibrils soluble fraction. **h**, MSA-amplified fibril soluble fraction. Similar results were observed in more than three independent experiments examining amplified fibrils derived from caudate samples for three cases of LBD and three cases of MSA.

Our previous studies demonstrated seeding activity in both soluble and insoluble fractions from MSA tissue, whereas seeding activity is detected only in the insoluble fractions from LBD tissue (Fig. 2e-f)^47^. We observed similar properties for amplified fibrils. Seeding activity was observed in the 100,000x g supernatant for MSA amplified fibrils but not for PD amplified fibrils. (Fig. 2g-h).

### 2D classification of single-particle cryo-EM data indicates a two-protofilament structure with pseudo-2_1_ helical screw symmetry

We collected single-particle cryo-EM data on a sample of the isotopically-labeled LBD fibrils used for SSNMR analysis. Images from several 2D class averages obtained after 2D classification in RELION are consistent with a two-protofilament structure with pseudo-2_1_ helical screw symmetry (Extended Data Fig. 2). All 2D classes displayed very low twist, which limits the ability to refine a high-resolution 3D density map with helical reconstruction^48-51^. We also observed 2D classes constituting a low percentage of the total particles that appear consistent with single protofilament fibril structure, based on comparison of the diameter and density pattern to two protofilament classes, indicating that single protofilament fibrils are present as a minority population in the amplified fibril preparation.

### Solid-state NMR Spectroscopy of LBD fibrils

We leveraged the templated fibril amplification methodology to produce uniformly ^13^C,^15^N and ^13^C,^2^H,^15^N labeled LBD fibrils. The ^13^C-^13^C 2D correlation spectrum display a set of very sharp peaks that can be individually assigned to resonances within the core of the fibril (residues ∼34 to ∼100) (Fig. 3a). This spectrum suggests a highly ordered fibril core and a disordered N- and C-terminus, consistent with previous studies of fibrils and of Asyn in particular. We extended the resonance assignment of the non-amyloid beta component of plaque (NAC) core using ^13^C, ^2^H, ^15^N labeled fibrils. These fibrils were uniformly triply labeled but were amplified in 100% ^1^H_2_O (H_2_O) buffer, allowing for the protonation of exchangeable sites uniformly through the fibril. After amplification, fibrils were repeatedly washed with 100% ^2^H_2_O (D_2_O) buffer. This procedure produces fibrils that remain highly protonated at the amide of the backbone of the fibril core but are perdeuterated in the N- and C-terminus, making these resonances invisible to ^1^H detection experiments. In contrast, the backbone amide ^1^H atoms within the fibril core are protected from chemical exchange with the D_2_O and so the amide SSNMR signals are well resolved. Thus fibrillization in H_2_O and washing with D_2_O has the effect of increasing the resolution and sensitivity of the core residues for assignment and structural experiments. Fig. 3b shows a representative 2D projection of a (Fig. 3b, left) CO-N-H 3D and (Fig. 3b, right) CA-N-H experiments with the backbone assignments of ^13^C, ^2^H, ^15^N indicated. In addition to confirming the backbone ^13^C and ^15^N assignments determined for the uCN sample, additional assignments were made for residues K43, Q62, E83, G84, A85, and G86, which were better resolved in the ^1^H-detected experiments than in the ^13^C-detected versions. We estimated the relative protonation quantity and the rigidity of the backbone resonances by measuring the intensity of the cross peak of the 3D CA-N-H data (Fig. 3c). The data provided by the ^13^C and ^1^H detection experiments were leveraged to yield dihedral restraints (Fig. 3d) and predicted random coil index order parameter (RCI S^2^), revealing LBD fibrils form highly ordered beta-sheet like structure involving E34 – K45 and V63 to V95. Notably, there is a high correlation between the RCI order parameter and the measured intensity pattern of the 3D CA-N-H peaks (Fig. 3c). Additional details regarding SSNMR pulse sequences, spectra and assignments are outlined in Supplementary Notes and Extended Data Fig. 3-9.

**Fig. 3:**
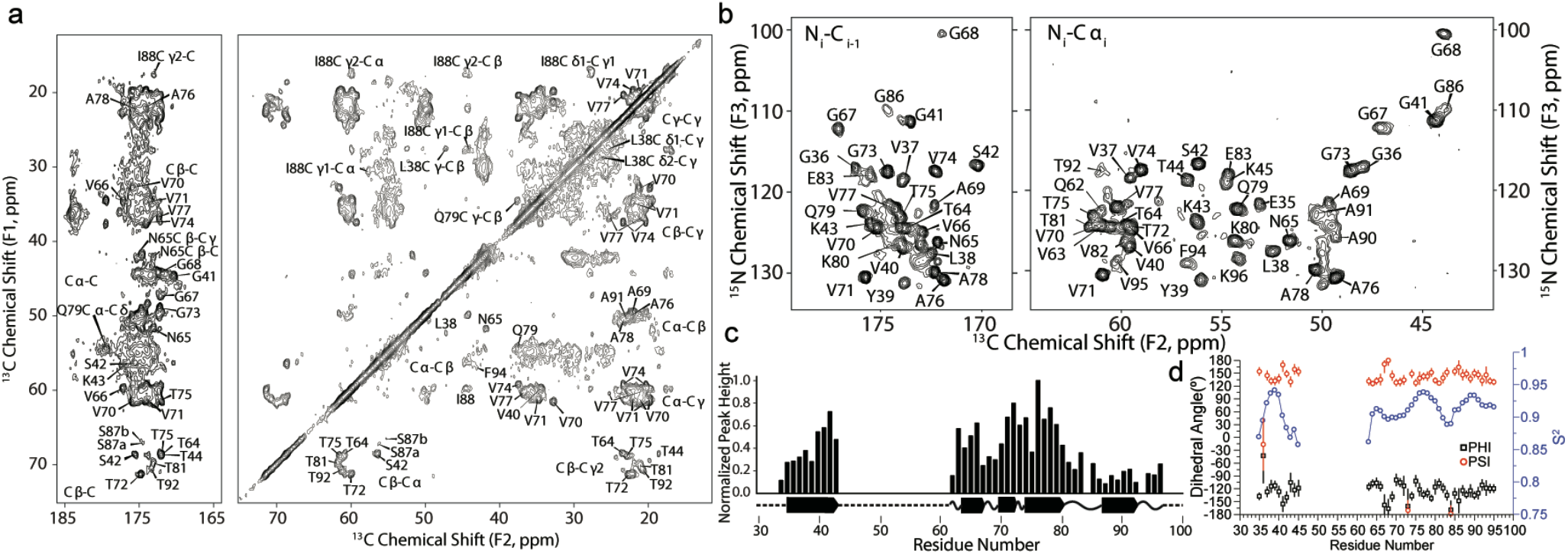
SSNMR assignments, long range contacts, and MPL measurements showing 2 protofilaments are the dominant population of fibrils. **a**, ^13^C-^13^C correlation of ^13^C-^15^N labeled LBD fibril showing carbonyl (left panel) and aliphatic (right panel) with unambiguous assignments labeled on the spectrum. Data was acquired at 750 MHz ^1^H frequency with 16.667 kHz magic-angle spinning and sample temperature 10±5 °C with 75 ms DARR mixing. **b**, 2D projections of a (left) CO-N-H 3D and (right) CA-N-H experiment with backbone assignments of a ^13^C, ^2^H, ^15^N labeled sample. Data in (b) were acquired at 750 MHz ^1^H frequency with 33.333 kHz magic-angle spinning, using a 6 ms ^15^N-^13^C cross polarization. 3Ds were collected using non-uniform sampling followed by reconstruction using SMILE prior to Fourier transformation. **c**, Peak heights are plotted for unambiguous assignments from a uniformly sampled CANH 3D spectrum collected on the uniformly ^13^C,^15^N-labeled sample. Bars are normalized to the peak height for residue A76. Predicted secondary structure from TALOS-N is plotted below the peak height plot for the LBD tissue-seeded sample. **d**, TALOS-N derived dihedral Phi and Psi angles reveal a beta-sheet structure. Predicted RCI chemical shift ordered parameter is plotted in blue showing highly ordered (R^2^ >0.9) regions between G36 and G41, T64 and T81, and A85 to V95.

We further measured long-range correlations to generate a structural model of the LBD amplified fibrils. Fig. 4a shows assignment strips of long-range unambiguous correlations between regions of the fibril that may be assigned using a ^13^C-^13^C correlation experiment with 12 ms of PAR mixing (see also Extended Data Fig. 4). L38CA shows correlations to A78CA/CB; V70CB shows correlations to N65CA/CB; G73CA shows a key correlation to F94CD, which was then used to network assign the F94CB. The 2D also reveals cross peaks from S87CB to either A78CA or A91CA, as well as multiple cross peaks between the I88 methyl groups and the Q79 and V77 spin systems. To disambiguate some of the other regions of spectral overlap, PAR mixing was incorporated into 3D pulse sequences. The 3D ^13^C-^13^C-^13^C pulse sequence (Extended Data Fig. 5) used RFDR mixing separating the two indirect dimensions so that the first residue involved in the PAR transfer can be identified based on two ^13^C frequencies instead of just one. Additionally, T92CB and T72CB are clearly resolved in the 3D ^13^C-^13^C-^13^C and independently exhibit distinct cross peaks with F94CD (Fig. 4b). Additional correlations from the 3D ^13^C-^13^C-^13^C include (Extended Data Fig. 6) cross peaks between the G41CA and A69CA/CB and V70CB; N65CB and V70CA/CB/CG; G68CA to S42CA/CB and G41C; T75CB to T92CB; and T92CB to V74C’/CA/CB and G73C’. We additionally used ^1^H detection methodologies to gather long range correlations. Fig. 4c shows contacts identified within a hCAhhNH experiment showing close contact between L38 with A76 and V77 and G41 with A69.

**Fig. 4:**
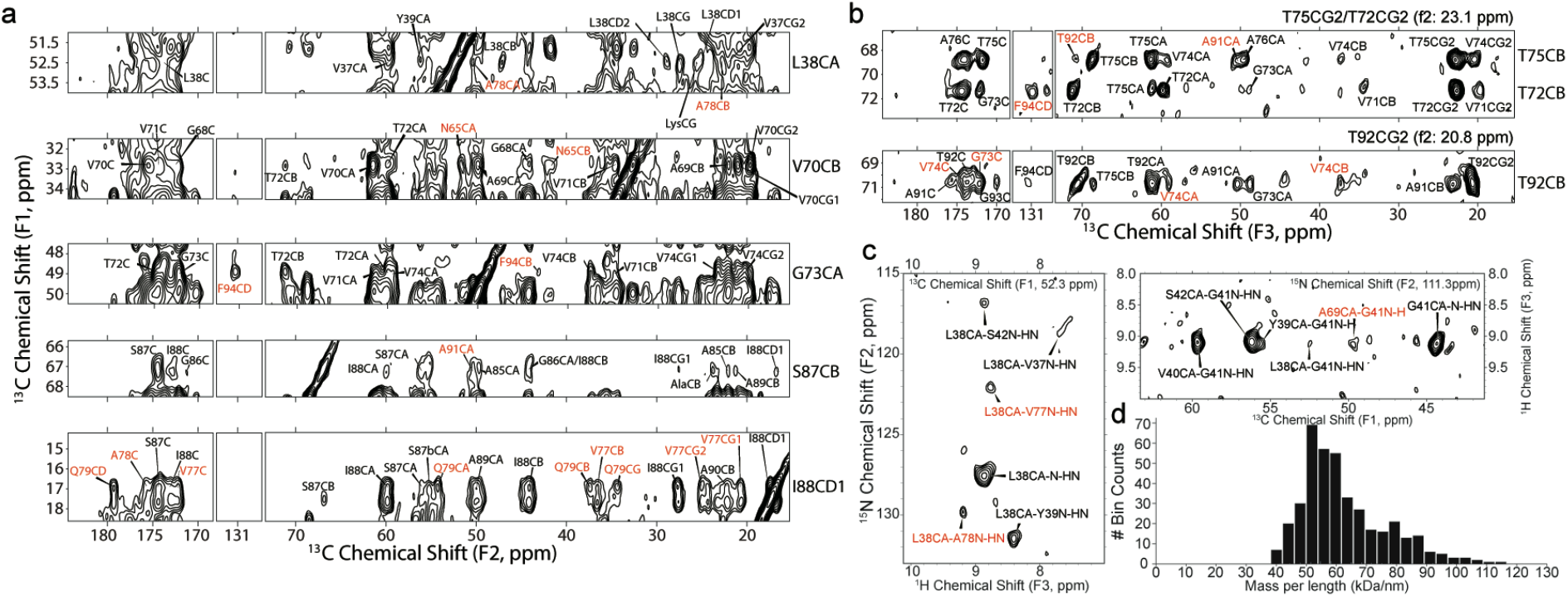
Long-range correlations determined by SSNMR. **a**, Long-range unambiguous correlations identified using 12 ms of PAR mixing between L38 V70, G73, S87, I88 found uniquely resolved within a ^13^C-^13^C 2D correlation experiment. **b**, 2D strip plots from a 3D ^13^C-^13^C-^13^C correlation spectrum showing correlations from three threonine (T72, T75, T92) residues that are disambiguated in F1 and F2 relative to the 2D spectrum. **c**, Long-range contacts identified within a hCAhhNH experiment showing close contact between L38 with A76 and V77 and G41 with A69. **d**, Mass per unit length histogram of the LBD fibril measured with dark-field, unstained TB-TEM micrograph.

Mass-per-length (MPL) is an invaluable constraint on the supramolecular organization of amyloid fibrils that can be measured experimentally by TEM. Dark-field transmission electron microscopy (TEM) of unstained, amplified LBD Asyn fibrils from the exact same batch as those used to prepare the uCN SSNMR sample was used to measure an MPL of 60±14 kDa/nm (Fig. 4d). With a molecular weight of the uCN labeled monomer of 15.244±2 kDa as determined by MALDI-TOF MS (data not shown), the symmetry of the fibril must consist of two molecules of Asyn per 4.8 Å β-sheet spacing. This observation is also consistent with the appearance of a two-protofilament structure with pseudo-2_1_ helical screw symmetry in 2D class averages obtained from single particle cryo-EM data (Extended Data Fig. 2). For the calculation reported here, it was assumed that the fibril is parallel and in-register, and composed of two symmetric protofilaments, which is consistent with the observation of primarily one set of peaks in the SSNMR spectra.

### Atomic model of amplified LBD Asyn fibrils

In the calculation of the structure of amplified LBD Asyn fibrils, we utilized the PASD protocol^52,53^ to perform automated assignment of long-range correlations for generation of distance restraints in Xplor-NIH^54^. Xplor-NIH’s strict symmetry facility was used to simplify the calculation by representing the fibril using a single protomer subunit, replicated by rotational and translational symmetry to form a 10-subunit fibril in which a pair of five subunit two-fold symmetric protofilaments interact laterally. Violation statistics for high-likelihood restraints in the context of the calculated structure bundles at each stage of PASD are summarized in Extended Data Fig. 8a-b. Given the resonance list for the ^13^C-detected nuclei, the initial PASD assignment process resulted in 1099 cross-peaks from data collected on the uCN and diluted samples for which only non-short-range (between residues separated by 3 or more residues in primary sequence) assignments were possible. 262 ± 7, or 24% of these peaks contained assignments which were satisfied by the final ensemble of 11 structures. This initial procedure resulted in an average ambiguity of 20 ± 22 peak assignments per peak. Following the network analysis stage where reinforcing assignments are given higher weight, and including a filter to remove any peaks with a protomer ambiguity greater than two, the fraction of peaks satisfying the final bundle was at 31%, and the overall ambiguity dropped to 3 ± 2. Three rounds of PASD structure calculations (pass 2, pass 3, and pass 4) simultaneously removed nearly all the violating restraints and gradually added back the satisfied restraints. A hybrid manual/PASD pass5 structure calculation-based refinement was then performed, where the original peaks were reassigned based on structures generated using the pass4 peak lists, increased the number of satisfied restraints to 183 ± 2.

The structural model in Fig. 5a is a cartoon representation of the lowest energy structure produced from the series of calculations of two protofilaments, showing residues A30–N103. The 10 lowest energy structures of the LBD fibril are represented in Fig. 5b. Structural features include a tight interface between the two protofilaments, a core composed exclusively of hydrophobic and polar residues, and charged residues lining the periphery. Notably, the residues that make up the hydrophobic core of the fibril (V71 to V82) have been described before as essential to fibril formation^55^. Additionally, a likely salt bridge between E35 and K80 was not observed directly but was added as a modeling restraint during the final refinement calculation based on a combination of the FS-REDOR result and the observation that these two residues were consistently in close proximity in the converged structures, even without the salt bridge restraint. The top ten structures ranked according to lowest total energy in Xplor-NIH are overlaid by all heavy atoms in Fig. 5b. Residues 35–46 and 62–96 of these structures converged to a backbone RMSD of 1.2 Å and a heavy atom RMSD of 1.7 Å, a resolution that allows many individual side chains to be resolved.

**Fig. 5:**
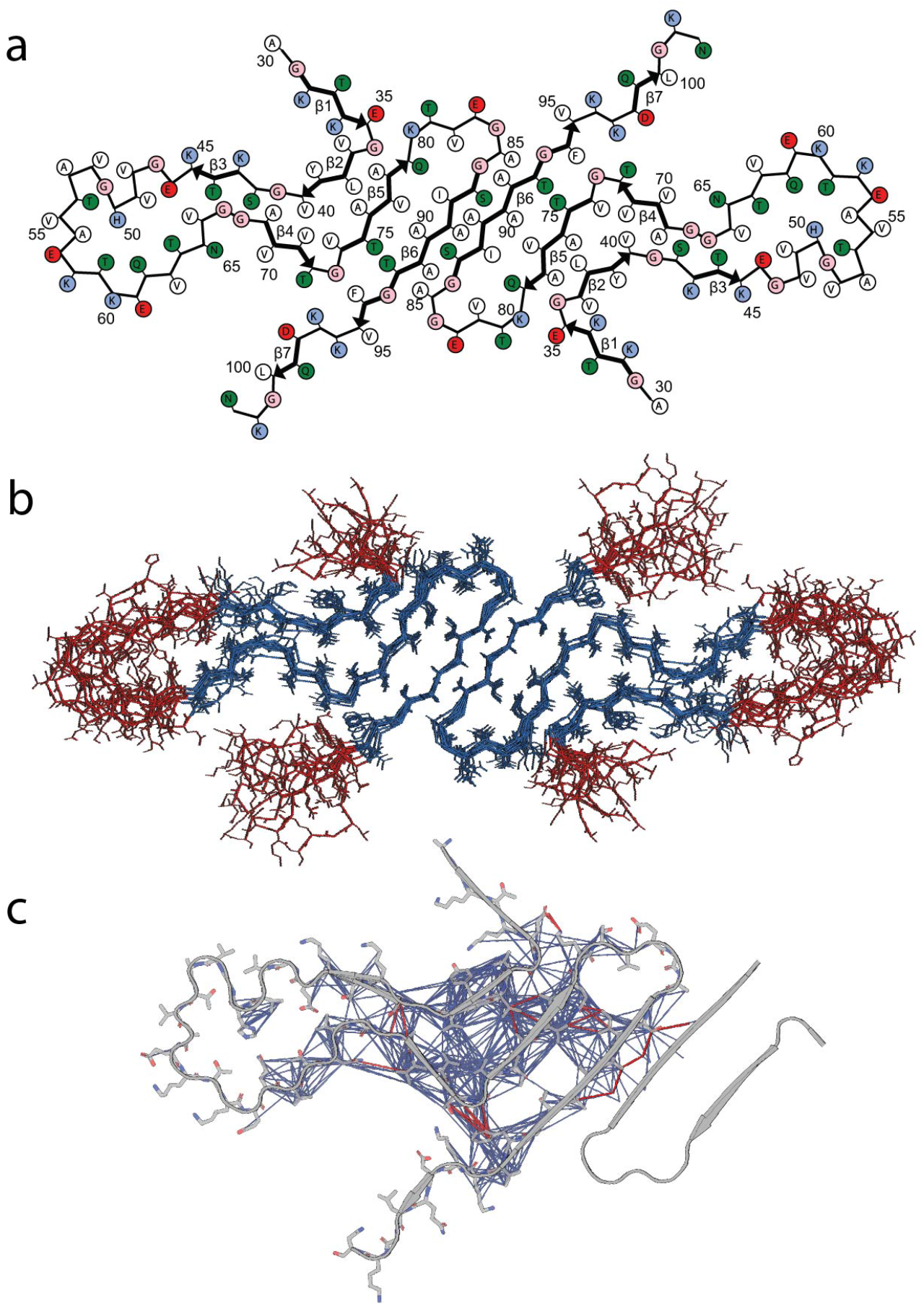
Structure of LBD fibrils. **a**, Cartoon representation of the LBD fibrils determined via SSNMR studies. **b**, 10 lowest energy structures of LBD fibril determined. Blue represent well-ordered regions with large amount of intermolecular contacts; red represent relatively disordered regions with few intermolecular contacts detectable using CP based polarization techniques. **c**, Inter-residue contacts identified from SSNMR CP based experiments. Magenta lines are intramolecular contacts manually identified within spectra, blue lines are correlations identified utilizing the PASD algorithm for probabilistic assignments.

We observe several notable structural features of the final determined structure. L38 forms a steric zipper with A76, V77, and A78 (Fig. 5a). G41 and S42 pack tightly against G68 and A69 (Fig. 5a). Correlations between the F94 aromatic carbons with T72 and G73, as well as the strong cross peaks between V71 with V74 and T75 with T92 are demonstrated. The interface between the protofilaments was particularly notable but also a challenge to assign manually because several short- and medium-range correlations are present (Fig. 5c). The PASD calculations result in convergence to the anti-parallel inter-filament beta-strand conformation. The resulting structure clarifies that S87 and A78 are part of the strong network of peaks between I88 with V77 and Q79, and that additional correlations from S87 to A91 on the other protofilament. We utilized an unrestrained molecular dynamics (MD) simulation to analyze the thermodynamic stability of the structural model determined by SSNMR and find that the core residues are qualitatively unchanged after 200 ns of equilibrium MD (Extended Data Fig. 9).

### SPARK analysis validates key amino acid residues required for LBD fibril growth and stability

We measured growth rates of amplified LBD fibrils in the presence of WT and mutant Asyn monomer, utilizing FlAsH dye to detect close association of two or more bicysteine tagged Asyn monomers upon incorporation into fibrils^39^. In this approach, which we refer to as Synuclein Polymorph Analysis by Relative Kinetics (SPARK), we examine relative differences in fibril growth rates that distinguish different fibril forms. We initially compared the effect of familial PD-associated Asyn mutations on growth rates for in vitro assembled (Tris) fibrils, LBD amplified fibrils and MSA amplified fibrils (Fig. 6a). For in vitro fibrils, we observed very inefficient growth, relative to WT, with the A53T and E46K mutations, but no effect on growth for H50Q^39^. MSA amplified fibrils are distinguished by an increased growth rate for A53T and minimal growth rate for E46K, consistent with a previous study demonstrating that MSA brain tissue extracts cannot seed aggregation of E46K Asyn in 293 cells^56^ further support for the fidelity of fibril amplification. LBD fibrils in contrast have much less pronounced relative differences in growth rates for these mutant monomers, yet still display a characteristic pattern for the four LBD cases we examined.

**Fig. 6:**
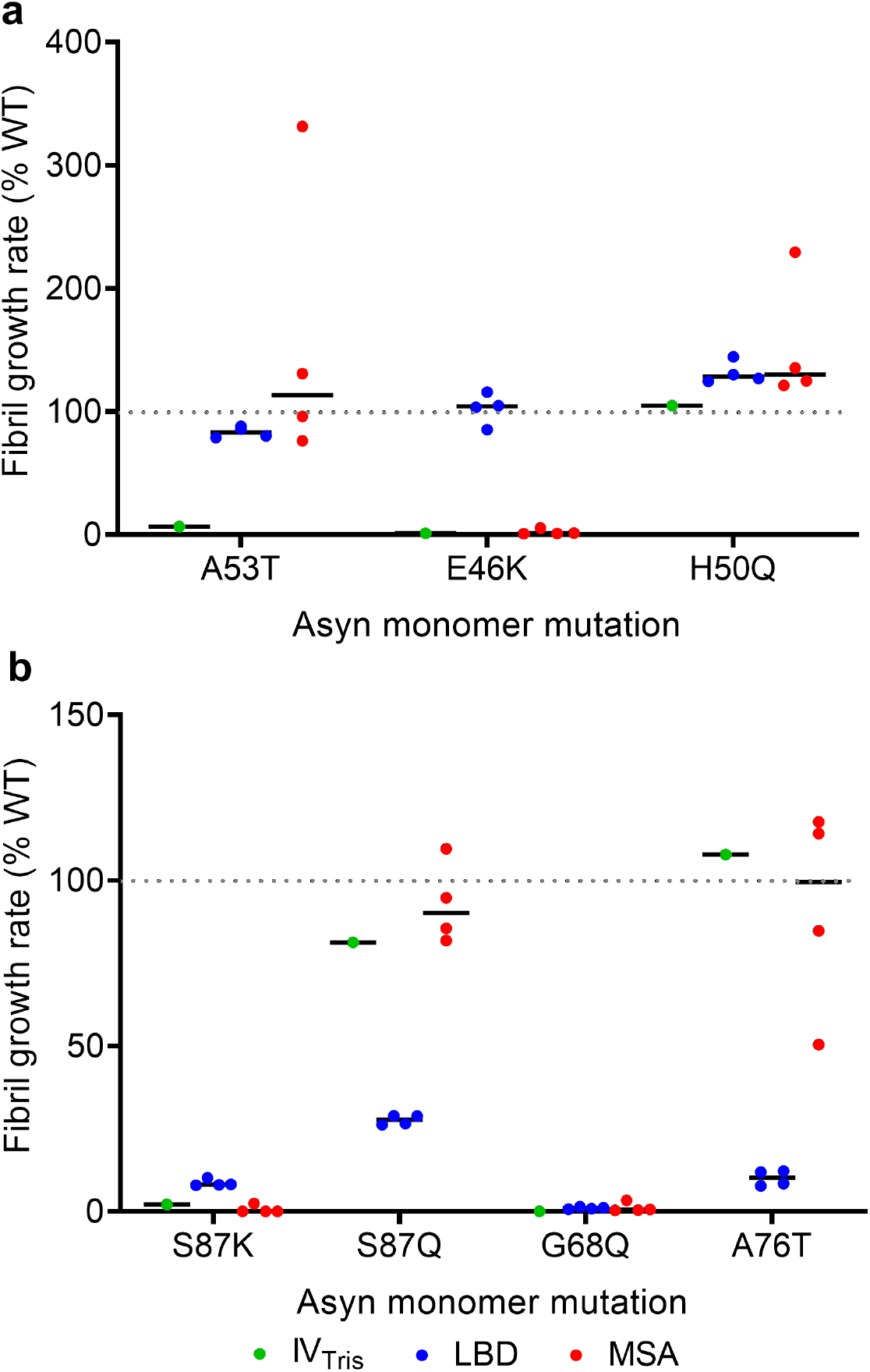
Fibril growth rates for in vitro assembled fibrils and amplified fibrils in the presence of mutant Asyn monomer. **a**, WT-normalized growth rates in the presence of Asyn monomer with familial PD mutations A53T, E46K and H50Q. **b**, WT-normalized growth rates in the presence of Asyn monomer with S87K, S87Q, G68Q and A76T mutations. The results show that the A53T mutation distinguishes in-vitro (Tris buffer) fibrils from the LBD and MSA amplified fibril forms and that the E46K-mutation distinguishes MSA amplified fibrils from LBD amplified fibrils. The S87Q and A76T mutations distinguish LBD amplified fibrils from in-vitro and MSA amplified fibril forms.

Based on the SSNMR structural model of LBD fibrils, we identified several residues central to fibril structure and examined the effect of these mutations on fibril growth. The S87K mutation dramatically reduces fibril growth for all three polymorphs (Fig. 6b). In contrast, a conservative S87Q mutation has little effect on in vitro and MSA fibrils, but substantially reduces LBD fibril growth. Similarly, an A76T mutation greatly inhibits LBD fibril growth, but has minimal effects on in vitro and MSA fibrils. Finally, G68Q dramatically reduces growth of all three polymorphs. Thus, we identified two mutations (S87K and G68Q) that critically alter growth and stability across multiple polymorphs, but also identified two mutations (S87Q and A76T) that are uniquely important for LBD fibril growth, further validating the structure. We attribute the large effect at S87 to the presence of the protofilament interface in the majority of LBD Asyn fibrils.

### Comparison of structural features among Asyn fibrils

The ordered core region of the amplified LBD Asyn fibrils is highly similar to the recently reported cryo-EM structure of fibrils extracted from postmortem PD and LBD brain tissue (Fig. 7) (PDB 8A9L^38^). Strands comprising residues 84-95 and 67-81 in the SSNMR and cryo-EM models overlay with root-mean-square deviation (r.m.s.d) values of the main-chain atoms of 0.73 Å and 1.0 Å, respectively. Moreover, these residue ranges have comparable numbers of satisfied SSNMR distance restraints (Fig. 7d-e). In the ‘hinge’ region (residues 47-61), there are very few SSNMR assignments and restraints. Likewise, the cryo-EM model reported higher B-factors in this same region. The N-terminal strand comprising residues 36-45 is more divergent between the two models, with a higher r.m.s.d. value (3.0 Å). Both models contain proteinaceous entities packed against the predominantly hydrophobic residues 85-93. In the SSNMR model, this is the unique interface formed between two protofilaments with pseudo-2_1_ helical screw symmetry. For the cryo-EM model, which comprises a single protofilament, this is the unidentified peptide termed island B. The cryo-EM model also contains an additional unidentified island A peptide packed against residues 50-55. In the SSNMR model, this residue range contains too few assignments and restraints to ascertain potential peptide interactions. The fold of LBD fibrils differs substantially from MSA and in vitro assembled Asyn fibrils. However, one consistent feature shared by LBD, MSA and in vitro conformers is the presence of an L-shaped motif formed by hydrophobic residues 69-79 (Extended Data Fig. 10), which is consistently buried between two other hydrophobic strands to form a three-layered structure.

**Fig. 7:**
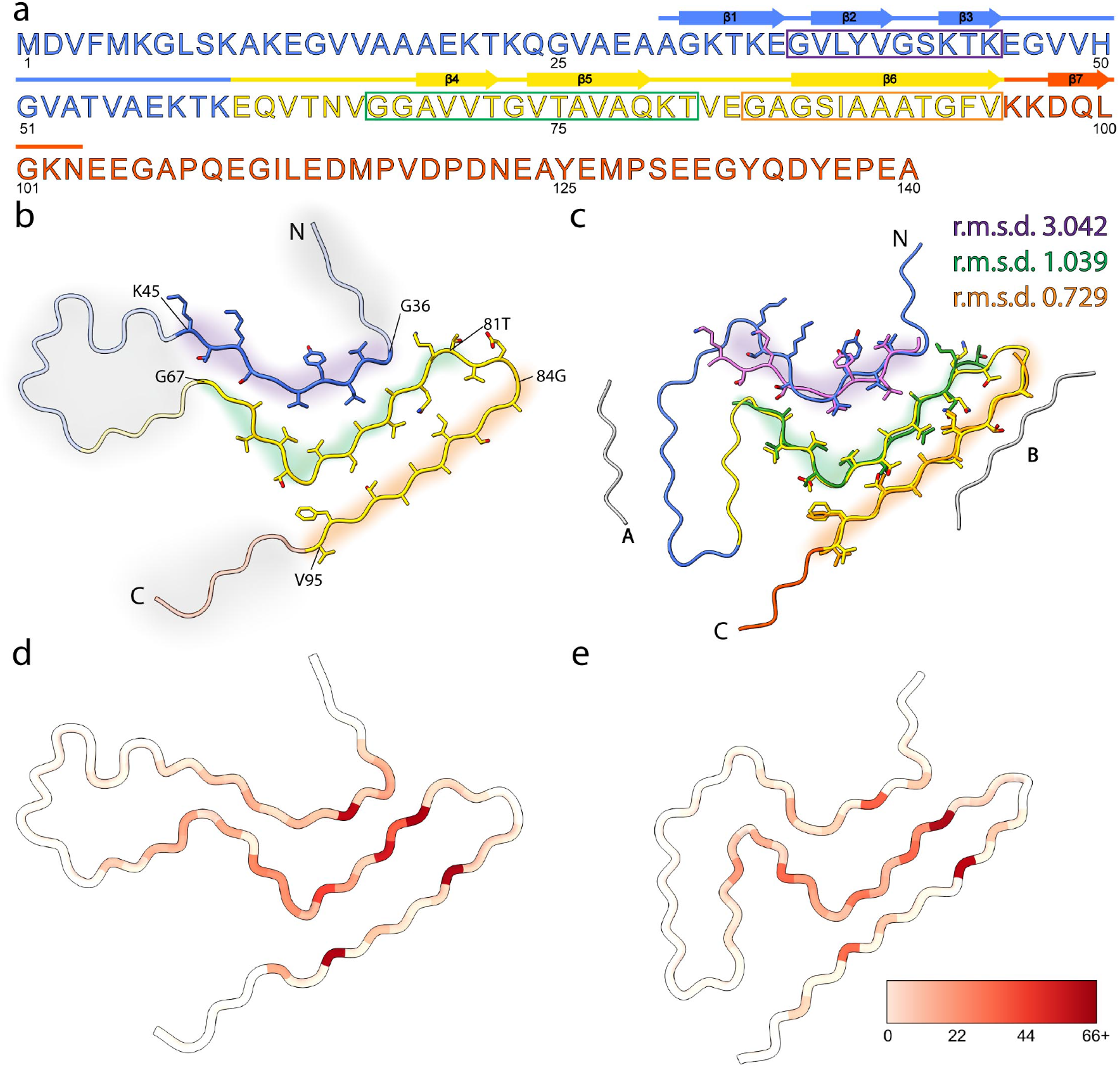
ssNMR structure of amplified Asyn resembles cryo-EM structure of extracted filaments. **a**, Primary sequence of human Asyn, with beta-strands annotated with arrows. The amphipathic N-terminal region, hydrophobic non-amyloid beta-component of plaque (NAC) region, and acidic C-terminal region are colored blue, yellow, and red, respectively. The purple, green, and orange rectangles denote the residue ranges used for subsequent comparison studies. **b**, Atomic model of SSNMR LBD Asyn core structure with labeled N- and C-termini (PDB 8FPT). Areas of low confidence are depicted with transparent cartoons with grey highlights. Residue ranges modeled with high confidence are depicted as cartoons with sidechains as sticks and are highlighted in purple, green, and orange. **c**, Atomic model of cryo-EM LBD Asyn core structure (PDB 8A9L), colored as in **b**. The unidentified proteinaceous islands are in grey. High-confident residue ranges of the SSNMR structure are overlayed in purple, green, and orange, and root-mean-square deviations (r.m.s.d) of the main-chain atoms were calculated. **d**, Number of satisfied NMR-derived interatomic distances mapped onto the lowest-energy structure using Xplor-NIH calculations on a per-residue basis using initial assignments from PASD. The color scale extends from white, indicating few or no NMR-derived distances were observed for the residue, to red, indicating many NMR-derived distances were observed for the residue. **e**, Number of satisfied NMR-derived distances mapped onto the cryo-EM Asyncore structure (PDB 8A9L).

## Discussion

We developed a new method to amplify Asyn fibrils from LBD postmortem tissue, enabling high-resolution analysis of fibril structure. MPL measurements with TEM, combined with 2D classification results from single-particle cryo-EM, indicate a two protofilament structure with pseudo-2_1_ helical screw symmetry. Several hundred SSNMR distance and dihedral restraints were used to develop a structural model based on backbone dihedral angle determination with TALOS-N^57^ and Xplor-NIH^54^ simulated annealing calculations. The fold of the two protofilament structural model is highly similar to the fold in the single protofilament structural model derived from single particle cryo-EM studies of LBD fibrils with high twist. These new results substantially extend the structural characterization of LBD Asyn fibrils by providing insight into the structure of fibrils with low twist that are not amenable to high resolution structure analysis with cryo-EM. They also indicate the existence of both two protofilament and single protofilament fibrils in LBD tissue.

There are several possible explanations for the identification of two protofilament fibrils in this study and single protofilament fibrils in the recent cryo-EM study^38^. In 2D classification of single-article cryo-EM data for amplified fibrils, we do observe minor classes of smaller diameter fibrils with a density pattern consistent with a single protofilament when compared to the predominant two-protofilament classes. The amplification protocol utilizes fibrils seeds obtained in a 1% Triton insoluble fraction, which is significantly different from the insoluble fraction obtained after extraction in 2% sarkosyl for the reported cryo-EM study. In contrast to extraction protocols utilizing sarkosyl, an ionic detergent with high solubilizing strength, we do not observe fibrils that are clearly isolated from the large amount of amorphous tissue-derived material after extraction in Triton, a weaker nonionic detergent. Thus, we were not able to directly assess diameter and structure of the fibril seeds used for amplification. It is possible that extraction with sarkosyl converts two protofilament fibrils to single protofilament fibrils, whereas a combination of single and two protofilament fibrils persists after extraction with Triton. It is also possible that our amplification protocol favors the accumulation of two protofilament fibrils over single protofilament fibrils, based on faster growth rates and/or higher fragmentation rates for two protofilament fibrils. Finally, it is possible that the sonication step in the amplification protocol promotes conversion of single protofilament fibrils to two protofilament fibrils by increasing the frequency of collision events. In situ studies of Asyn fibril structure in postmortem tissue, although technically challenging, could provide more definitive information on the presence and relative abundance of the two classes of fibrils.

By substantially expanding the availability of LBD fibril quantities beyond what can be obtained from frozen postmortem tissue samples, the amplified fibril preparations reported here provide new avenues for studies of disease mechanisms and therapeutics. For example, LBD amplified fibrils facilitate studies of Asyn fibril growth, as we demonstrate in the SPARK analysis. Amplified fibrils can also be used to seed LBD Asyn fibril accumulation in cell culture models and animal models, where the LBD fold is likely to be maintained in accumulating fibrils, a prediction that can ultimately be verified by structure analysis of fibrils derived model systems. The use of amplified fibrils in vitro or in model systems can facilitate the development of imaging agents by enabling in vitro and in vivo measurements of binding properties for Asyn ligands. Finally, studies in vitro and in model systems can facilitate the development of candidate therapeutics capable of inhibiting fibril accumulation. Other amplification methods, utilizing tissue extracts or cerebrospinal fluid as starting material, have produced Asyn fibrils with structures that diverge from the structures determined for tissue-derived fibrils in MSA and LBD, indicating that some aspects of the amplification protocol described here, such as fragmentation and incubation conditions, are important for templating structure during amplification^58-60^.

The fold of LBD Asyn fibrils is distinct from MSA Asyn fibrils and from all in vitro assembled polymorphs reported to date. LBD fibril structure appears to be highly similar among LBD cases, with similarity in the cryo-EM density maps obtained for six LBD cases in the study by Yang et al^38^. We have observed a high degree of similarity in SSNMR spectra for three other LBD cases in addition to the one reported here (data not shown). However, further studies are needed to determine whether subtle structural variations exist among LBD cases and also whether structural heterogeneity occurs within individual cases. Only 25% of particles with high twist were amenable to cryo-EM analysis and cryo-EM density maps were derived from only a subset of the particles with high twist^38^. Structural variations, if present, may relate to the spectrum of clinical features observed in LBD, such as the variable timing of dementia onset. They may also relate to genetic and environmental risk factors identified for PD/DLB. For example, different genetic and environmental factors may give rise to differences in metabolites, post-translational modifications and interacting proteins that drive variations on a set of core structural elements.

SPARK analysis validates and extends the structure analysis for LBD fibrils. Using SSNMR structural data, we identified four amino acid residues likely to be central to fibril structure, and validated that these residues are critical for LBD fibril growth and stability. Two of the amino acid residues are uniquely important for LBD fibrils when compared to in vitro and MSA amplified fibrils, providing additional insight into unique structural features of LBD fibrils. These differences in fibril growth rates likely reflect structural constraints specific to each fibril form, where a mutation of a specific amino acid residue has the potential to alter kinetics based on interactions with neighboring beta strands within a single monomeric unit or alternatively at the protofilament interface. Analysis of additional residues could extend this kinetic characterization of fibril structure. SPARK analysis could also be utilized to probe the significance of structural heterogeneity, if identified within individual LBD autopsy cases, or to analyze the significance of structural variations if observed across different LBD cases.

Further investigation of Asyn fibril structure in LBD can include analysis of fibrils obtained with additional extraction and amplification methods, with the goal of improved understanding of the conditions that give rise to single protofilament and two protofilament fibril forms, as well as twisted fibrils. Understanding conditions responsible for generating structural features specific to LBD fibrils may provide insight into disease mechanisms in LBD. Joint refinement of structural models based on SSNMR and cryo-EM data could enhance the structural information available to guide the development of imaging ligands and therapeutics, which will also be aided by analysis of additional LBD cases.

## Methods

### Demographics and clinical information of participants

The Movement Disorders Center Neuropathology Core, Washington University, St. Louis, MO, provided clinically and neuropathologically well-characterized postmortem frozen brain tissue (Supplementary Table 5). LBD1 was selected amplified for SSNMR structural characterization.

### Asyn monomer production (natural abundance)

Natural abundance recombinant Asyn protein was prepared using methods described previously. BL21(DE3)RIL bacterial cells were transformed with the pRK172 plasmid containing the WT Asyn construct^39^. The natural abundance Asyn protein was purified by heat denaturation and precipitation of bacterial proteins, followed by ion-exchange chromatography. Purified monomeric Asyn was dialyzed and stored at −80 °C. For use in amplified fibril production, the purified monomer was filtered through Amicon ultra-4, 50k (Catalog UFC805024, Millipore) cutoff filter to remove any preformed Asyn aggregates. The 50k-filtered Asyn monomer was stored at 4 °C until use in the amplification process. Micro-BCA assay was used to determine the protein concentration of the Asyn monomer.

### Preparation of Insoluble fraction seeds from LBD, MSA and control postmortem tissue

The protocol to sequentially extract human postmortem brain tissue was adapted from our previous publication^61^. Briefly, gray matter dissected from tissue was sequentially homogenized in four buffers (3 ml/g wet weight of tissue) using Kimble Chase Konte™ dounce tissue grinders (KT885300-0002). In the first step, 300mg of dissected grey matter tissue was homogenized using 20 strokes of Pestle A in High Salt (HS) buffer (50 mM Tris-HCl pH 7.5, 750 mM NaCl, 5 mM EDTA plus 0.1% (v/v) Sigma P2714 Protease Inhibitor (PI) cocktail). The homogenate was centrifuged at 100,000 ×g for 20 min at 4 °C and the pellet was homogenized in the next buffer using 20 strokes of Pestle B. Extractions using Pestle B were performed in HS buffer with 1% Triton X-100 with PI, then HS buffer with 1% Triton X-100 and 1 M sucrose, and with 50 mM Tris-HCl, pH 7.4 buffer. In the final centrifugation, the resulting pellet was resuspended in 50 mM Tris-HCl, pH 7.4 buffer (3 ml/g wet weight of tissue). The aliquots of insoluble fraction were stored at −80 °C until use. Similar extraction protocol was followed for LBD, MSA and control cases.

### Preparation of LBD, MSA and control postmortem soluble tissue fraction

Dissected gray matter of tissue (100-300 mg) was prepared by homogenization of dissected gray matter in 50 mM Tris, pH 7.4 (3 ml/g wet weight of tissue) using 20 strokes of Pestle A followed by 20 strokes of Pestle B dounce tissue homogenizers. The aliquots of unfractionated homogenates were stored at −80 °C until use. A 100 µl aliquot of this homogenate was spun down at 100,000 xg and the supernatant used as the soluble fraction in cell culture experiments.

### Amplification of LBD fibrils from LBD insoluble fraction seeds

We amplified LBD fibrils from gray matter dissected from caudate brain region. Insoluble fraction (10 µl) containing 3.3 µg wet wt. of tissue was bought to a final volume of 30 µl by addition of 20 mM Tris-HCl, pH 8.0 plus 100 mM NaCl buffer (fibril buffer) in a 1.7 ml microcentrifuge tube. This insoluble fraction was sonicated for 2 min at amplitude 50 in a bath sonicator (Qsonica model Q700) with a cup horn (5.5 inch) attachment at 4 °C. To the sonicated seeds, 1.5 µl of 2 % Triton X-100 was added to recover the sonicated tissue derived Asyn fibril seeds. To this mixture, 50k Amicon ultra filtered Asyn monomer was added to a final concentration of 2 mg/ml in a final volume of 100 µl. This mixture underwent quiescent incubation at 37 °C for 3 days, completing the 1^st^ cycle of sonication plus incubation. After the first cycle, the mixture was sonicated at 1 min at amplitude 50, and then an additional 300 µl of 2 mg/ml Asyn monomer was added. The mixture underwent quiescent incubation at 37 °C for 2 days (2^nd^ cycle). Then, sonication for 1 min at amplitude 50 and quiescent incubation for 2 days was repeated for the third cycle, followed by sonication for 1 min at amplitude 50 and quiescent incubation for 3 days for the 4^th^ cycle. At the end of 4^th^ cycle, LBD amplified fibrils were stored at 4 °C until use. The typical scheme of incubations was 3-2-2-3 days for 4 cycles.

Further expansion of the LBD amplified fibrils was performed by centrifuging 60 µl of the 4^th^ cycle LBD amplified fibrils at 21,000 xg for 15 min at 4 °C. The pellet was resupended in 100 µl of fibril buffer and sonicated for 1 min at amplified 50. To this mixture, Asyn monomer was added to a final concentration of 2 mg/ml in a final volume of 400 µl in fibril buffer. This mix was quiescently incubated at 37 °C for 2 days (5^th^ cycle). At the end of 5^th^ cycle, samples were centrifuged at 21,000 xg for 15 min at 4 °C and the top 300 µl of spent Asyn monomer was moved to a separate tube. The pellet was resuspended by trituration and sonicated for 1 min at amplified 50. After sonication, the previously removed monomer was added back. Next, an additional 2.5 mg/ml of Asyn monomer was added to bring the total volume to 800 µl. This mixture was incubated at 37 °C for 2 days to complete 6^th^ round of incubation. The increased monomer concentration (2.5 mg/ml instead of 2 mg/ml) was calculated based on the average decrease in free Asyn monomer due to its incorporation into amplified fibrils. The 6^th^ cycle fibrils were stored at 4 °C until use. In parallel, insoluble fraction from control tissue samples were also amplified under the similar conditions used for LBD amplified fibrils.

### Production of isotopically labeled Asyn monomer

Expression of uniform [^13^C, ^15^N] labeled wild-type a-synuclein was carried out in *E. coli* BL21(DE3)/pET28a-AS in modified Studier medium M^62^. The labeling medium contained 3.3 g/L [^13^C]glucose, 3 g/L [^15^N]ammonium chloride, 11 mL/L [^13^C, ^15^N]Bioexpress (Cambridge Isotope Laboratories, Inc., Tewksbury, MA), 1 mL/L BME vitamins (Sigma), and 90 µg/mL kanamycin. After a preliminary growth in medium containing natural abundance (NA) isotopes, the cells were transferred to the labeling medium at 37 ºC to an OD_600_ of 1.2, at which point the temperature was reduced to 25 ºC and protein expression induced with 0.5 mM isopropyl β-D-1-thiogalactopyranoside (IPTG) and grown for 15 h to a final OD_600_ of 4.1 and harvested.

For [2-^13^C]glycerol, uniform ^15^N sample diluted to 25% in NA monomer, the labeled monomer was expressed according to the protocol of Tuttle et al^16^. A preliminary culture was grown in one volume of Studier medium M containing 2g/L ammonium chloride, 2 g/L glycerol, 1 mL/L BME vitamins, and 90 µg/mL kanamycin to an OD_600_ of 2.0. The cells were gently pelleted and resuspended in 0.5 volume of Studier medium M containing 2g/L [^15^N]ammonium chloride, 4 g/L [2-^13^C]glycerol, 1 g/L [^13^C]sodium carbonate, 1 mL/L BME vitamins, and 90 µg/mL kanamycin. To maximize aeration, the culture was aliquoted as 150 ml per baffled 2-L flask, and grown at 25 ºC with shaking at 250 rpm. After an hour the OD_600_ was 4.4, and expression was induced with 0.5 mM IPTG. The culture was harvested 15 h later with an OD_600_ of 5.5.

For the uniform [^2^H, ^13^C, ^15^N] labeling, a frozen stock was prepared beforehand by highly expressing cells of BL21(DE3)/pET28a-AS using the “double colony selection” method of Sivashanmugam et al^63^. This stock was used to streak an LB plate containing 70% D_2_O and 40 µg/ml kanamycin which was grown overnight. The colonies were then used to inoculate LB-kanamycin in 70% D_2_O which was grown at 37 ºC to an OD_600_ of 3.6. The cells were gently pelleted and resuspended in an equal volume of 99% D_2_O SHD medium (modified from the Sivashanmugam et al. optimized High cell-Density medium^63^), containing 50 mM Na_2_HPO_4_, 25 mM KH_2_PO_4_, 10 mM NaCl, 5 mM MgSO_4_, 0.2 mM CaCl_2_, 8 g/L [^2^H, ^13^C]glucose, 1 g/L [^15^N]ammonium chloride, 10 mL/L [^2^H, ^13^C, ^15^N]Bioexpress, 2.5 mL/L BME vitamins, 2.5 mL/L Studier trace metals, 90 µg/mL kanamycin, and was adjusted to pD 8.0 with NaOD ^64^. Growth was continued with 40 ml culture per baffled 250 ml flask at 25 ºC and 250 rpm shaking, to promote aeration. At an OD_600_ of 4.3, expression was induced with 0.5 mM IPTG. The culture was harvested after 15 h with a final OD_600_ of 10.6.

Protein purification was done as described previously^65^. Briefly, cells were lysed chemically in the presence of Turbonuclease (Sigma) to digest nucleic acids. Purification began with a heat denaturation of the cleared lysate, followed by ammonium sulfate precipitation^66^. The resolubilized protein was bound to QFF anion exchange resin (GE Healthcare Life Sciences, Marlborough, MA) and eluted using a linear gradient of 0.2–0.6 M NaCl. Fractions containing Asyn monomer, which eluted at about 0.3 M NaCl, were pooled, concentrated, and run over a 26/60 Sephacryl S-200 HR gel filtration column (GE Healthcare Life Sciences) equilibrated in 50 mM Tris-HCl, 100 mM NaCl, pH 8 buffer. Fractions were pooled, concentrated to ∼20 mg/ml a-synuclein, and dialyzed at 4 ºC into 10 mM Tris-HCl pH 7.6, 50 mM NaCl, 1 mM DTT, and stored at a concentration of ∼14 mg/mL at −80 ºC until use. Yields were 95 mg purified AS protein/L growth medium for the uniform [^13^C, ^15^N] labeled monomer, 288 mg/L for the uniform [^2^H, ^13^C, ^15^N] labeled monomer, and 217 mg/L for the [2-^13^C]glycerol, ^15^N labeled monomer.

### Amplification of Isotopically labeled LBD-amplified fibrils for SSNMR studies

SSNMR experiments relied on large quantities of amplified fibrils. We selected postmortem case LBD1 for SSNMR characterization. Multiple replication tubes of LBD1 amplification were setup using uCN and uCDN Asyn monomers. The protocol to generate LBD amplified fibrils was similar to that described above, except that 50k Amicon ultra filtered isotopically labeled Asyn monomer was used. In addition, the amplification reactions were supplemented by control fraction preparation derived from E.Coli transformed with an empty expression vector. This control fraction was purified with the same protocol as the natural abundance Asyn monomer. Amplified fibrils were harvested by ultracentrifugation, washed with either H_2_O or D_2_O, dried under a gentle stream of nitrogen gas, packed into 1.6 mm rotors (Revolution NMR, LLC) and hydrated to approximately 40% with either H_2_O or D_2_O, as described^67^.

### Amplification of MSA fibrils from MSA insoluble fraction seeds

We produced MSA amplified fibrils from caudate brain region, matching the brain selected for LBD amplification. The protocol to generate MSA amplified fibrils was similar to the LBD amplified fibrils until the 4^th^ cycle. Further expansion was performed by bringing up 60 µl of 4^th^ cycle MSA amplified fibrils to 100 µl and sonicating the mixture for 1 min at amplitude 50. To this mixture, Asyn monomer was added to a final concentration of 2 mg/ml in a final volume of 400 µl in fibril buffer. This mixture was quiescently incubated at 37 °C for 2 days (5^th^ cycle). To expand further, the 400 µl of 5^th^ cycle MSA amplified fibrils were sonicated for 1 min at amplified 50 and additional 2.1 mg/ml of Asyn monomer was added to bring the total volume to 800 µl. This mix was incubated at 37 °C for 2 days to complete 6^th^ cycle of incubation. The increased monomer concentration (2.1 mg/ml instead of 2 mg/ml) was calculated based on the loss of free Asyn monomer due to its incorporation into amplified fibrils. The 6^th^ cycle fibrils were stored at 4 °C until use. In parallel, insoluble fraction from control tissue samples were also amplified under the similar conditions used for LBD amplified fibrils.

### Radioligand quantification of amplified fibrils

Radioligand binding assay was used to estimate the specificity and quantify the growth of LBD amplified fibrils at the end of amplification process. A modified version of homologous competition binding assay with radioligand [^3^H]-BF2846 was utilized to assess the fibrils at the end of the 4^th^ and 6^th^ cycles. We assumed equal affinity and binding site density of the radiotracer to LBD and control amplified fibrils. A fixed volume of amplified fibrils (LBD and control) were added to either 300 nM cold BF2846 (made in DMSO) or with no cold (DMSO only) along with 2 nM of [^3^H]-BF2846. The total volume of reaction mixture was 150 µl in 30 mM Tris-HCl, pH 7.4 plus 0.1% BSA buffer. The mix was incubated for 2 h at 37 °C with shaking (1000 rpm) in an Incu-Mixer MP2 (Catalog H6002 Benchmark Scientific). Samples were transferred to Multiscreen FB Filter plates (Catalog MSFBN6B50) and washed three times with ice-cold 30 mM Tris-HCl, pH 7.4 plus 0.1% BSA buffer. The glass fiber filters containing fibril bound radioligand were removed and counted overnight in a PerkinElmer (1450-021) Trilux MicroBeta Liquid Scintillation counter using 150 µl of Optiphase Supermix cocktail (Perkin Elmer). All data points were done in triplicate. The displacement counts between the two different cold concentrations was converted to concentration of fibrils and used for comparison of specificity between disease and control amplifications.

### Concentration estimation of LBD and control amplified fibrils by Micro BCA assay

We utilized the micro-BCA assay to estimate the amount of fibrils. At the end of the 6^th^ cycle, amplified fibrils were centrifuged at 21,000 xg for 15 min at 4 °C to separate fibrils from monomer. The concentration of Asyn monomer in the supernatant was determined by the Micro BCA protein assay (Thermo Scientific Pierce Micro BCA kit, catalog no. 23235) according to the manufacturer’s instructions, using the manufacturer-supplied bovine serum albumin (BSA) for the standard curve. At the same time, total sample containing fibrils plus unincorporated monomer was also assessed. The measured decrease in Asyn monomer concentration between total and supernatant was used to determine the concentration of fibrils.

### Characterization of amplified fibrils via negative stain TEM

Negative staining of the different fibril conformers was performed by applying a given fibril preparation to ultrathin Carbon 300 mesh Gold grids (Catalog 01824G, Ted Pella). The grids were negatively glow discharged (13 mA, 45 sec) using GloQube glow discharge system (Model #025235 EMS). A 10 µl fibril sample at appropriate dilution was applied to the glow discharged grid for 5 minutes with carbon side facing the sample drop. Post-sample incubation, the grid was washed (6 times) with 50 µl of H_2_O, washed once with 50 µl of 0.75% uranyl formate and stained with 50 µl of 0.75% uranyl formate for 3 minutes. The grids were blotted using filter paper, leaving a small amount of stain to air dry on the grid surface. Grids were imaged on a JEOL 1400 TEM operating at 120kV to visualize negatively staining fibrils.

### Cryo-EM Grid Preparation and Imaging

Amplified Asyn fibril samples were prepared for cryo-EM imaging on Quantifoil holey carbon grids (R2/2 300 mesh copper) by plunge freezing using a Vitrobot Mark IV (ThermoFisher Scientific, Brno, CZ). Grids were plasma cleaned for 1 minute using H2/O2 plasma in a Gatan Solarus 950 (Gatan, Pleasanton, CA) prior to plunge freezing. The sample chamber of the Vitrobot was set to 4°C and 95% humidity. 3 µL of sample was applied to the grid surface. After 20 s of incubation, the grids were blotted for 2 s with a blot force of 4 and plunge frozen into liquid ethane. Single particle cryo-EM imaging was performed using a Cs-corrected ThermoFisher Titan Krios G3 cryo-electron microscope (ThermoFisher Scientific, Eindhoven, NL) operating at a 300 kV accelerating voltage. EPU (ThermoFisher Scientific, Brno, CZ) was used for image acquisition using a Falcon IV direct electron detector (ThermoFisher Scientific, Eindhoven, NL) at a magnification of 75,000x which corresponds to a 0.9 Å pixel size. Movies were acquired for 8.38 s and used 48 total frames. The total dose was 53.04 electrons per Å2 and the total dose per frame was 1.105 electrons per Å2 per frame. The defocus range was −1 to −2.4 µm.

### Cryo-EM Image Processing

Cryo-EM data was processed using helical reconstruction methods in RELION^48^. Movies were motion-corrected and dose-weighted using MOTIONCOR2^68^. Motion-corrected micrographs were then to estimate the contrast transfer function (CTF) using GCTF^69^. Asyn filaments were first picked manually to generate 2D templates via 2D Classification in cryoSPARC. Once low-resolution templates were obtained, they were used to train the Filament Tracer helical auto-picking feature. The final selection of particles picked by Filament Tracer were extracted using a box size of 280 pixels with a 0.9 Å pixel size. Initial particles were reduced to a final set of ∼63k after several rounds of 2D classification in cryoSPARC. This particle set was then exported into RELION, via csparc2star.py, for an additional round of 2D classification.

### Cell culture experiments

To assess the fidelity of amplification, we utilized previously described HEK293T “biosensor” cell line stably expressing α-syn (A53T)-CFP/YFP fusion proteins^47^. Glass coverslips were coated with poly-d-lysine hydrobromide (Sigma) overnight at room temperature, then washed 3 times with sterile water and allowed to air-dry for 2 h. Biosensor cells were plated in a 24 well plate at 75,000 cells per well and grown for 1 day in 400 µl of cell culture media. Biosensor cells were then treated with amplified fibrils, insoluble fraction or the soluble fractions derived from the amplified fibrils and the postmortem tissue. Seeding samples were diluted in Opti-MEM (Gibco) to a final volume of 50 µl, then, sonicated for 1 min at amplitude 50 in the bath sonicator. To this, 3 µl of Lipofectamine 3000 in 50 µl Opti-MEM was added and the mixture incubated for 30 min at room temperature for complex formation. This 100 µl of fibrils plus lipofectamine mixture was added dropwise to 400 µl of the culture medium in each well of a 24 well plate. For insoluble and soluble fraction derived from tissue, we added 5 µl per well. For LBD amplified fibrils samples, concentrations between 26 nM to 1 nM were used. For MSA amplified fibril samples, concentrations between 1.5-0.1 nM was used. After 72 h, biosensor cells were fixed with 4% paraformaldehyde (EMS) for 15 min and washed with Dulbecco’s PBS three times. Coverslips were then mounted on glass slides using Fluoromount-G containing DAPI (Southern Biotechnology). Coverslips were imaged on a Nikon Eclipse TE2000-U fluorescence microscope using a Nikon Plan Fluor ×40/0.75 and a 10x objective. Natural abundance Asyn monomer amplified samples (LBD3, LBD4, LBD5, MSA 1, MSA2 and MSA4) were selected for the cell culture experiments.

### Solid-state NMR spectroscopy

Magic-angle spinning (MAS) SSNMR experiments were performed at magnetic field of 11.7 T (500 MHz ^1^H frequency) or 17.6 T (750 MHz ^1^H frequency) using Varian NMR (Walnut Creek, CA) VNMRS spectrometers. Spinning was controlled with a Varian MAS controller to 11,111 ± 30 Hz or 22,222 ± 15 Hz (500 MHz ^1^H frequency) and 16,667 ± 15 Hz or 33,333 ± 30 Hz (17.6 T), with two exceptions indicated in Supplementary Table 6. Sample temperature during SSNMR data collection was 10±5 °C. The 11.7 T magnet was equipped with a 1.6 mm HCDN T3 probe (Varian), with pulse widths of about 1.8 µs for ^1^H and ^13^C, and 3.2 µs for ^15^N. The 17.6 T magnet was equipped with a HXYZ T3 probe (Varian) tuned to HCN triple resonance mode with pulse widths of 1.9 µs for ^1^H, 2.6 µs for ^13^C, and 3.0 µs for ^15^N. All experiments utilized ^1^H-^13^C or ^1^H-^15^N tangent ramped CP^70^ and ∼100 kHz SPINAL-64 decoupling during evolution and acquisition periods^71,72^. Where applicable, SPECIFIC CP was used for ^15^N-^13^Cα and ^15^N-^13^C’ transfers^73^; ^13^C-^13^C homonuclear mixing was performed using DARR^74^, RFDR^75,76^, or PAR^77^; and water suppression was done using MISSISSIPPI^78^. FS-REDOR was performed according to Jaroniec et al^79^. The reference spectrum was acquired with a 630 µs Gaussian pi pulse centered on resonance with the Glu CDs, while the REDOR dephasing was performed using 9 µs square pi pulses placed every half rotor period on the ^15^N channel, except for a 540 µs Gaussian pi pulse in the middle of the dephasing period placed on resonance with the Lys NZs. Chemical shifts were externally referenced to the downfield peak of adamantane at 40.48 ppm^80^. NUS schedules using biased exponential sampling were prepared using the nus-tool application in NMRbox^81^. Data conversion and processing was done with NMRPipe ^82^. NUS data was first expanded with the nusExpand.tcl script in NMRPipe, converted, and processed using the built-in SMILE reconstruction function^83^. Peak picking and chemical shift assignments were performed using SPARKY 3.

### Mass-per Unit Length measurements with Electron Microscopy

TEM grids were prepared using a 1 min oxygen plasma cleaning treatment (Harrick Plasma Cleaner PDC-32G, at low power). One 10 µl droplet of Asyn fibril suspension and three 10 µl droplets of ultrapure water for each grid were pipetted onto a clean sheet of Parafilm. Freshly plasma-cleaned TEM grids were inverted onto a sample droplet and rested for 60 s. Excess sample solution was blotted away with filter paper, and grids were placed onto each of the three droplets of water and blotted again in quick succession, to rinse away excess salt. Grids were then rested on a droplet of tobacco mosaic virus (TMV) suspension, prepared by diluting a stock solution to 0.12 mg/mL. TMV was used to calibrate electron density in each image^84-86^. Imaging was done on a JEOL 2100 Cryo-TEM using an electron accelerating voltage of 80 kV. Micrographs were collected in the tilt-beam geometry using the third objective aperture, an exposure time of 3–5 s. Short, non-overlapping segments of TMV and Asyn fibrils in each image were then selected using the helixboxer function of EMAN2^87^ and exported to the MpUL-multi program for quantification of MPL statistics^88^. Segments were only chosen from TMV and Asyn fibrils when they were distinguishable in both tilt-beam and accompanying bright-field micrographs.

### Xplor-NIH Structure Calculations

Automated structure calculation and refinement was done using the PASD protocol^52,53^ in Xplor-NIH^54^. Four SSNMR spectra were used for assignment of long-range correlations: (1) 2D ^13^C-^13^C with 12 ms PAR mixing collected on the uCN sample; (2) 3D ^13^C-^13^C-^13^C with 1.9 ms RFDR then 12 ms PAR mixing collected on the uCN sample; (3) 3D ^15^N-^13^Ca-^13^CX with 12 ms PAR mixing collected on the uCN sample; and (4) 2D ^13^C-^13^C with DARR mixing collected on the diluted sample. Peaks were picked manually for 2Ds or using restricted peak picking for 3Ds in SPARKY 3 then filtered against the resonance list to remove any peaks without at least one assignment possibility using an in-house Python script. These lists were used directly as input for the automated generation of peak assignments using PASD.

Calculations were performed using the strict symmetry facility in Xplor-NIH^54^. Based on the model of two protofilaments that are symmetrically arranged, only a single protomer subunit was explicitly simulated, while four additional subunits were generated by translation along the z-axis by the centroid position of the first protomer, and five were generated by rotation of 180 degrees about the z-axis followed by translation. This approach simplified restraint generation because the only interactions that were critical to simulate were those between the first protomer (A1) and one generated by translational symmetry (B1), and those between the first protomer (A1) and one in-plane generated by rotational symmetry (A2). A restraint with an atom-atom ambiguity of one in terms of the number of possible resonance assignments therefore had a total ambiguity of five (intramolecular, A1-A1; intermolecular, A1-B1, B1-A1, A1-A2, A2-A1). The intramolecular B1-B1 and A2-A2 were identical by symmetry and therefore were not included as assignment options. Likewise, the intermolecular/interfilament B1-A2 and A2-B1 were also excluded.

To match cross-peak positions to assigned chemical shift resonances, the following tolerances were used: 0.25–0.35 ppm for indirect ^13^C dimensions, 0.35 ppm for indirect ^15^N, and 0.20–0.25 ppm for direct dimensions. Automated chemical shift correction was not performed. Distances for all three PAR mixing data sets were binned as strong (up to 4.5 Å), medium (up to 6.0 Å), weak (up to 7.0 Å), and very weak (up to 8.5 Å) and the DARR were binned as strong (upto 4.5 Å), medium (up to 5.5 Å), weak (up to 6.5 Å), and very weak (up to 7.5 Å). Long-range assignment options for the isotopically diluted sample data were assumed to be exclusively intramolecular, whereas all possibilities were included as ambiguous for the uCN and uCDN sample data. After initial matching of peak lists to the resonance list, the network filter was applied. In addition to assigning likelihoods of 0 or 1 to peak assignments based on the network filter analysis, peaks with intraresidue assignment possibilities or peaks with at least one peak assignment that was short-range (differed in primary sequence by less than 3 residues), or peaks with more than two possible atom-atom peak assignments were removed. This reduced file sizes and computation time by significantly reducing the number of interactions that need to be computed.

The initial coordinates for each structure calculation were those of a geometry-optimized protomer with backbone dihedral angles satisfying the TALOS-N restraint list, and truncated to residues 30– 103, excluding the disordered N- and C-terminal regions that are not in the fibril core. Calculation of 500 structures for each pass (pass 2, pass 3, and pass4) was performed according to the protocol in Supplementary table 8. In addition to the PASD peak lists, a few manually assigned distance restraints were included in all structure calculations to help guide simulated annealing. These included backbone hydrogen bond registry restraints based on non-exchanging ^1^H signals observed in the uCDN sample, seven restraints that were hypothesized to be exclusively intrafilament based on some preliminary structure calculations, and three restraints that were hypothesized to be exclusively interfilament using similar logic. Statistics were collected on the 200 lowest energy structures for each pass. Peak assignment likelihood reanalysis following each PASD structure calculation pass was done using the 20 best-converged structures from the low energy fraction. Pass 2 and pass 3 were run using only peaks with an atom-atom peak assignment ambiguity of 1 or 2. Likelihood reanalysis based on the pass 3 structures was done using the pass 1 peak lists, which re-introduced the originally higher ambiguity peaks, now at significantly lower ambiguity. Following pass 4, the high-likelihood restraints (≥ 0.9) were converted into Xplor-NIH format and used in a pass 5 standard simulated annealing calculation employing 192 structures. The 19 structures with lowest energy were used for an additional likelihood reanalysis and as input into a structure-dependent PASD refinement (pass 4), which was otherwise identical to pass 3 and pass 4. A final single-assignment restraint list was generated using a few iterative rounds of regular simulated annealing calculations starting with the Xplor-NIH restraint lists based on pass 5, manual adjustment to the restraint lists based on study of the violation statistics, and automated peak assignment likelihood reanalysis. At this stage, medium range peak assignments (including restraints between neighboring adjacent in sequence from the initial matching) were also incorporated to help with optimization of local side chain orientations. The final refinement calculation also included an intrafilament salt bridge modeling restraint between residues E35 and K80 based on the detection from FS-REDOR (see Fig. Extended Data Fig. 8) that the structure contains at least one salt-bridge between a Glu CD and a Lys NZ and the fact that this salt bridge was observed in the majority of converged structures even without this modeling restraint being present.

### Assessing Structural Agreement from PASD Assignments

NMR peak lists were used to assess agreement of the structures with NMR assignments. Peak lists with associated NMR assignments were loaded into Xplor-NIH as PASD potentials and assignments that did not agree with observed distances in a provided reference structure to within 0.7 angstroms were given a likelihood of zero. All peaks that agreed were assigned a likelihood of one and the “to” and “from” atomic assignments for these peaks were extracted and compiled on a per-residue basis. The total assignments per residue are mapped onto Figure 7 (d,e) as heat maps.

### Molecular Dynamics (MD) Simulations and Analysis

An all-atom MD simulation of a two-fold symmetric Asyn fibril (2×20-mer) was performed using NAMD 3.0^89^. The CHARMM36m protein force field was applied to the fibrils^90^. We simulated the fibril in an explicit water box of size 180 × 100 x 180 Å3 with an ion concentration of 0.15 M (NaCl solution) using an NPT ensemble with N = 306450, P = 1.01325 bar and T = 310 K. The system was subjected to Langevin dynamics with damping constant γ = 1.0 ps-1 and the Nosé-Hoover Langevin piston method^91,92^ was employed to maintain constant pressure. The MD integration time step was set to 2 fs. To evaluate long range electrostatic interactions, particle mesh Ewald^93^ was used in the presence of periodic boundary conditions. Nonbonded van der Waals (vdW) interactions were calculated with the Lennard-Jones (12, 6) potential with a cutoff distance of 12 Å. A smoothing function was introduced at 10 Å to gradually truncate the vdW potential energy at the cut-off distance. 14 Å cutoff distance was used to identify the atom pairs for vdW interaction. The system was first energetically minimized for 1000 steps by conjugate gradient method followed by a 10 ns equilibration of water and ions in the presence of positional restraints (force constant: 1 kcal/mol Å2) on Cα atoms of the fibril. The system was then simulated for 200 ns without any restraints. Simulation trajectories were collected every 50 ps.

### Synuclein Polymorph Analysis by Relative Kinetics (SPARK) assays

We developed a sensitive assay that uses the fluorescein arsenical dye FlAsH-EDT_2_ (Invitrogen TC-FlAsH™ in-cell tetracysteine tag detection kit, catalog no. T34561) to measure fibril growth rates in the presence of C2-WT and mutant C2-Asyn monomer. The SPARK assay was similar to FlAsH seed growth experiments demonstrated in our previous publication^39^. Amplified fibril samples at 1.5 µM concentration in fibril buffer plus 0.1% Triton X-100 were sonicated for 5 min at amplitude 50 in the cup horn sonicator (Qsonica) and then mixed with 0.5 mg/ml of C2-Asyn monomer (C2-WT-Asyn and mutant C2-Asyn) in a total volume of 25 µl. The seeds plus monomer mixture were quiescently incubated for 3 h at 37 °C in Corning Black 96-well plates (Fisher, catalog no. 07-200-762). After 3 h, FlAsH assay mixture consisting of 3.5 mm tris(2-carboxyethyl)phosphine, 1 mm EDT, 1 mm EDTA, 25 nm FlAsH-EDT_2_, and 200 mM Tris-HCl, pH 8.0 was added and the mixture incubated for additional 1 h at room temperature. FlAsH fluorescence was detected in a BioTek plate reader using a 485/20-nm excitation filter, a 528/20-nm emission filter, top 510-nm optical setting, and gain setting 100. Data was normalized to fibril growth rates with WT-C2-Asyn monomer. We selected 6^th^ cycle LBD and MSA amplified fibrils (case LBD1, LBD2, LBD3, LBD4, MSA1, MSA2, MSA3 and MSA4) for SPARK assays.

## Acknowledgements

Support for this work was provided by: grants from the Michael J. Fox Foundation; NIH grants NS110436, NS097799, and NS075321 from the National Institute of Neurological Disorders and Stroke and National Institute on Aging and P41GM136463 from the National Institute of General Medical Sciences. CDS was supported by the Intramural Research Programs of NIDDK and NHLBI at the National Institutes of Health. We thank Deborah Berthold for assistance with production of isotopically-labeled recombinant Asyn protein.

## Author contributions

Study design and organization: DD, AB, CR, PK; Acquisition of data: DD, AB, CB, JL, JO, MR, SS, JS; Analysis and interpretation of data: DD, AB, CB, KB, IG, MM, MR, ZS, SS, BS, JS, OW, QC, JF, CS, ET, CR, PK; Drafting of the manuscript: DD, AB, CB, KB, MR, CR, PK; Critical revision of the manuscript for important intellectual content: All authors.

## References

1. Forno, L.S. Neuropathology of Parkinson’s disease. J.Neuropathol.Exp.Neurol. 55, 259–272 (1996).

2. Appel-Cresswell, S. et al. Alpha-synuclein p.H50Q, a novel pathogenic mutation for Parkinson’s disease. Mov Disord 28, 811–3 (2013).

3. Golbe, L.I., Di Iorio, G., Bonavita, V., Miller, D.C. & Duvoisin, R.C. A large kindred with autosomal dominant Parkinson’s disease. Ann.Neurol. 27, 276–282 (1990).

4. Kruger, R. et al. Ala30Pro mutation in the gene encoding alpha-synuclein in Parkinson’s disease. Nat.Genet. 18, 106–108 (1998).

5. Lesage, S. et al. G51D alpha-synuclein mutation causes a novel parkinsonian-pyramidal syndrome. Ann Neurol 73, 459–71 (2013).

6. Polymeropoulos, M.H. et al. Mutation in the alpha-synuclein gene identified in families with Parkinson’s disease. Science 276, 2045–2047 (1997).

7. Proukakis, C. et al. A novel alpha-synuclein missense mutation in Parkinson disease. Neurology 80, 1062–4 (2013).

8. Singleton, A.B. et al. alpha-Synuclein locus triplication causes Parkinson’s disease. Science 302, 841 (2003).

9. Zarranz, J.J. et al. The new mutation, E46K, of alpha-synuclein causes Parkinson and Lewy body dementia. Ann.Neurol. 55, 164–173 (2004).

10. Compta, Y. et al. Lewy- and Alzheimer-type pathologies in Parkinson’s disease dementia: which is more important? Brain 134, 1493–1505 (2011).

11. Hely, M.A., Reid, W.G., Adena, M.A., Halliday, G.M. & Morris, J.G. The Sydney multicenter study of Parkinson’s disease: the inevitability of dementia at 20 years. Mov Disord. 23, 837–844 (2008).

12. Hurtig, H.I. et al. Alpha-synuclein cortical Lewy bodies correlate with dementia in Parkinson’s disease. Neurology 54, 1916–1921 (2000).

13. Brundin, P., Dave, K.D. & Kordower, J.H. Therapeutic approaches to target alpha-synuclein pathology. Exp Neurol 298, 225–235 (2017).

14. Bousset, L. et al. Structural and functional characterization of two alpha-synuclein strains. Nat Commun 4, 2575 (2013).

15. Gath, J. et al. Unlike twins: an NMR comparison of two alpha-synuclein polymorphs featuring different toxicity. PLoS One 9, e90659 (2014).

16. Tuttle, M.D. et al. Solid-state NMR structure of a pathogenic fibril of full-length human alpha-synuclein. Nat Struct Mol Biol 23, 409–15 (2016).

17. Peelaerts, W. & Baekelandt, V. ɑ-Synuclein strains and the variable pathologies of synucleinopathies. J Neurochem 139, 256–274 (2016).

18. Melki, R. Role of Different Alpha-Synuclein Strains in Synucleinopathies, Similarities with other Neurodegenerative Diseases. Journal of Parkinson’s Disease 5, 217–227 (2015).

19. Lemkau, L.R. et al. Site-specific perturbations of alpha-synuclein fibril structure by the Parkinson’s disease associated mutations A53T and E46K. PLoS One 8, e49750 (2013).

20. Gath, J. et al. Yet another polymorph of alpha-synuclein: solid-state sequential assignments. Biomol NMR Assign 8, 395–404 (2014).

21. Verasdonck, J. et al. Further exploration of the conformational space of alpha-synuclein fibrils: solid-state NMR assignment of a high-pH polymorph. Biomol NMR Assign 10, 5–12 (2016).

22. Guerrero-Ferreira, R. et al. Cryo-EM structure of alpha-synuclein fibrils. bioRxiv (2018).

23. Guerrero-Ferreira, R. et al. Two new polymorphic structures of human full-length alpha-synuclein fibrils solved by cryo-electron microscopy. Elife 8(2019).

24. Li, B. et al. Cryo-EM of full-length alpha-synuclein reveals fibril polymorphs with a common structural kernel. Nat Commun 9, 3609 (2018).

25. Sun, Y. et al. Cryo-EM structure of full-length alpha-synuclein amyloid fibril with Parkinson’s disease familial A53T mutation. Cell Res 30, 360–362 (2020).

26. Boyer, D.R. et al. Structures of fibrils formed by alpha-synuclein hereditary disease mutant H50Q reveal new polymorphs. Nat Struct Mol Biol 26, 1044–1052 (2019).

27. Boyer, D.R. et al. The alpha-synuclein hereditary mutation E46K unlocks a more stable, pathogenic fibril structure. Proc Natl Acad Sci U S A 117, 3592–3602 (2020).

28. Zhao, K. et al. Parkinson’s disease-related phosphorylation at Tyr39 rearranges alpha-synuclein amyloid fibril structure revealed by cryo-EM. Proc Natl Acad Sci U S A 117, 20305–20315 (2020).

29. Li, Y. et al. Amyloid fibril structure of α-synuclein determined by cryo-electron microscopy. Cell Res 28, 897–903 (2018).

30. Ni, X., McGlinchey, R.P., Jiang, J. & Lee, J.C. Structural Insights into α-Synuclein Fibril Polymorphism: Effects of Parkinson’s Disease-Related C-Terminal Truncations. J Mol Biol 431, 3913–3919 (2019).

31. McGlinchey, R.P., Ni, X., Shadish, J.A., Jiang, J. & Lee, J.C. The N terminus of α-synuclein dictates fibril formation. Proc Natl Acad Sci U S A 118(2021).

32. Sun, Y. et al. The hereditary mutation G51D unlocks a distinct fibril strain transmissible to wild-type α-synuclein. Nat Commun 12, 6252 (2021).

33. Long, H. et al. Wild-type α-synuclein inherits the structure and exacerbated neuropathology of E46K mutant fibril strain by cross-seeding. Proc Natl Acad Sci U S A 118(2021).

34. Zhao, K. et al. Parkinson’s disease associated mutation E46K of α-synuclein triggers the formation of a distinct fibril structure. Nat Commun 11, 2643 (2020).

35. Tao, Y. et al. Heparin induces α-synuclein to form new fibril polymorphs with attenuated neuropathology. Nat Commun 13, 4226 (2022).

36. Frieg, B. et al. The 3D structure of lipidic fibrils of α-synuclein. Nat Commun 13, 6810 (2022).

37. Schweighauser, M. et al. Structures of α-synuclein filaments from multiple system atrophy. Nature 585, 464–469 (2020).

38. Yang, Y. et al. Structures of alpha-synuclein filaments from human brains with Lewy pathology. Nature (2022).

39. Dhavale, D.D. et al. A sensitive assay reveals structural requirements for alpha-synuclein fibril growth. J Biol Chem 292, 9034–9050 (2017).

40. Ferrie, J.J. et al. Identification of a nanomolar affinity α-synuclein fibril imaging probe by ultra-high throughput in silico screening. Chemical Science 11, 12746–12754 (2020).

41. Hsieh, C.J. et al. Alpha Synuclein Fibrils Contain Multiple Binding Sites for Small Molecules. ACS Chem Neurosci 9, 2521–2527 (2018).

42. Bagchi, D.P. et al. Binding of the Radioligand SIL23 to alpha-Synuclein Fibrils in Parkinson Disease Brain Tissue Establishes Feasibility and Screening Approaches for Developing a Parkinson Disease Imaging Agent. PLoS One 8, e55031 (2013).

43. Barclay, A.M., Dhavale, D.D., Courtney, J.M., Kotzbauer, P.T. & Rienstra, C.M. Resonance assignments of an alpha-synuclein fibril prepared in Tris buffer at moderate ionic strength. Biomol NMR Assign 12, 195–199 (2018).

44. Chu, W. et al. Design, Synthesis, and Characterization of 3-(Benzylidene)indolin-2-one Derivatives as Ligands for alpha-Synuclein Fibrils. J Med Chem 58, 6002–17 (2015).

45. Holmes, B.B. et al. Heparan sulfate proteoglycans mediate internalization and propagation of specific proteopathic seeds. Proc Natl Acad Sci U S A 110, E3138–47 (2013).

46. Hsieh, C.J. et al. Chalcones and Five-Membered Heterocyclic Isosteres Bind to Alpha Synuclein Fibrils in Vitro. ACS Omega 3, 4486–4493 (2018).

47. Yamasaki, T.R. et al. Parkinson’s disease and multiple system atrophy have distinct alpha-synuclein seed characteristics. J Biol Chem 294, 1045–1058 (2019).

48. He, S. & Scheres, S.H.W. Helical reconstruction in RELION. J Struct Biol 198, 163–176 (2017).

49. Kimanius, D., Forsberg, B.O., Scheres, S.H. & Lindahl, E. Accelerated cryo-EM structure determination with parallelisation using GPUs in RELION-2. Elife 5(2016).

50. Scheres, S.H. RELION: implementation of a Bayesian approach to cryo-EM structure determination. J Struct Biol 180, 519–30 (2012).

51. Scheres, S.H. A Bayesian view on cryo-EM structure determination. J Mol Biol 415, 406–18 (2012).

52. Kuszewski, J. et al. Completely Automated, Highly Error-Tolerant Macromolecular Structure Determination from Multidimensional Nuclear Overhauser Enhancement Spectra and Chemical Shift Assignments. Journal of the American Chemical Society 126, 6258–6273 (2004).

53. Kuszewski, J.J., Thottungal, R.A., Clore, G.M. & Schwieters, C.D. Automated error-tolerant macromolecular structure determination from multidimensional nuclear Overhauser enhancement spectra and chemical shift assignments: improved robustness and performance of the PASD algorithm. J Biomol NMR 41, 221–39 (2008).

54. Schwieters, C.D., Bermejo, G.A. & Clore, G.M. Xplor-NIH for molecular structure determination from NMR and other data sources. Protein Sci 27, 26–40 (2018).

55. Giasson, B.I., Murray, I.V., Trojanowski, J.Q. & Lee, V.M. A hydrophobic stretch of 12 amino acid residues in the middle of alpha-synuclein is essential for filament assembly. J Biol Chem 276, 2380–6 (2001).

56. Woerman, A.L. et al. Familial Parkinson’s point mutation abolishes multiple system atrophy prion replication. Proc Natl Acad Sci U S A 115, 409–414 (2018).

57. Shen, Y. & Bax, A. Protein backbone and sidechain torsion angles predicted from NMR chemical shifts using artificial neural networks. J Biomol NMR 56, 227–41 (2013).

58. Burger, D., Fenyi, A., Bousset, L., Stahlberg, H. & Melki, R. Cryo-EM structure of alpha-synuclein fibrils amplified by PMCA from PD and MSA patient brains. bioRxiv, 2021.07.08.451588 (2021).

59. Fan, Y. et al. Different structures and pathologies of α-synuclein fibrils derived from preclinical and postmortem patients of Parkinson’s disease. bioRxiv, 2021.11.02.467019 (2021).

60. Frieg, B. et al. α-Synuclein polymorphism determines oligodendroglial dysfunction. bioRxiv, 2021.07.09.451731 (2021).

61. Miller, R.L. et al. Quantifying regional α -synuclein, amyloid β, and tau accumulation in lewy body dementia. Annals of Clinical and Translational Neurology 9, 106–121 (2022).

62. Studier, F.W. Protein production by auto-induction in high density shaking cultures. Protein Expr Purif 41, 207–34 (2005).

63. Sivashanmugam, A. et al. Practical protocols for production of very high yields of recombinant proteins using Escherichia coli. Protein Sci 18, 936–48 (2009).

64. Glasoe, P.K. & Long, F.A. USE OF GLASS ELECTRODES TO MEASURE ACIDITIES IN DEUTERIUM OXIDE1,2. The Journal of Physical Chemistry 64, 188–190 (1960).

65. Barclay, A.M., Dhavale, D.D., Courtney, J.M., Kotzbauer, P.T. & Rienstra, C.M. Resonance assignments of an α-synuclein fibril prepared in Tris buffer at moderate ionic strength. Biomol NMR Assign 12, 195–199 (2018).

66. Kloepper, K.D., Woods, W.S., Winter, K.A., George, J.M. & Rienstra, C.M. Preparation of alpha-synuclein fibrils for solid-state NMR: expression, purification, and incubation of wild-type and mutant forms. Protein Expr Purif 48, 112–7 (2006).

67. Tuttle, M.D., Courtney, J.M., Barclay, A.M. & Rienstra, C.M. Preparation of Amyloid Fibrils for Magic-Angle Spinning Solid-State NMR Spectroscopy. Methods Mol Biol 1345, 173–83 (2016).

68. Zheng, S.Q. et al. MotionCor2: anisotropic correction of beam-induced motion for improved cryo-electron microscopy. Nature Methods 14, 331–332 (2017).

69. Zhang, K. Gctf: Real-time CTF determination and correction. J Struct Biol 193, 1–12 (2016).

70. Metz, G., Wu, X.L. & Smith, S.O. Ramped-Amplitude Cross Polarization in Magic-Angle-Spinning NMR. Journal of Magnetic Resonance, Series A 110, 219–227 (1994).

71. Fung, B.M., Khitrin, A.K. & Ermolaev, K. An Improved Broadband Decoupling Sequence for Liquid Crystals and Solids. Journal of Magnetic Resonance 142, 97–101 (2000).

72. Comellas, G., Lopez, J.J., Nieuwkoop, A.J., Lemkau, L.R. & Rienstra, C.M. Straightforward, effective calibration of SPINAL-64 decoupling results in the enhancement of sensitivity and resolution of biomolecular solid-state NMR. Journal of Magnetic Resonance 209, 131–135 (2011).

73. Baldus, M., Petkova, A.T., Herzfeld, J. & Griffin, R.G. Cross polarization in the tilted frame: assignment and spectral simplification in heteronuclear spin systems. Molecular Physics 95, 1197–1207 (1998).

74. Takegoshi, K., Nakamura, S. & Terao, T. 13C–1H dipolar-assisted rotational resonance in magic-angle spinning NMR. Chemical Physics Letters 344, 631–637 (2001).

75. Bennett, A.E., Griffin, R.G., Ok, J.H. & Vega, S. Chemical shift correlation spectroscopy in rotating solids: Radio frequency-driven dipolar recoupling and longitudinal exchange. The Journal of Chemical Physics 96, 8624–8627 (1992).

76. Bennett, A.E. et al. Homonuclear radio frequency-driven recoupling in rotating solids. The Journal of Chemical Physics 108, 9463–9479 (1998).

77. De Paëpe, G., Lewandowski, J.R., Loquet, A., Böckmann, A. & Griffin, R.G. Proton assisted recoupling and protein structure determination. J Chem Phys 129, 245101 (2008).

78. Zhou, D.H. & Rienstra, C.M. High-performance solvent suppression for proton detected solid-state NMR. J Magn Reson 192, 167–72 (2008).

79. Jaroniec, C.P., Tounge, B.A., Herzfeld, J. & Griffin, R.G. Frequency Selective Heteronuclear Dipolar Recoupling in Rotating Solids: Accurate 13C−15N Distance Measurements in Uniformly 13C,15N-labeled Peptides. Journal of the American Chemical Society 123, 3507–3519 (2001).

80. Morcombe, C.R. & Zilm, K.W. Chemical shift referencing in MAS solid state NMR. Journal of Magnetic Resonance 162, 479–486 (2003).

81. Maciejewski, M.W. et al. NMRbox: A Resource for Biomolecular NMR Computation. Biophys J 112, 1529–1534 (2017).

82. Delaglio, F. et al. NMRPipe: a multidimensional spectral processing system based on UNIX pipes. J Biomol NMR 6, 277–93 (1995).

83. Ying, J., Delaglio, F., Torchia, D.A. & Bax, A. Sparse multidimensional iterative lineshape-enhanced (SMILE) reconstruction of both non-uniformly sampled and conventional NMR data. J Biomol NMR 68, 101–118 (2017).

84. Chen, B., Thurber, K.R., Shewmaker, F., Wickner, R.B. & Tycko, R. Measurement of amyloid fibril mass-per-length by tilted-beam transmission electron microscopy. Proc Natl Acad Sci U S A 106, 14339–44 (2009).

85. Komatsu, H., Feingold-Link, E., Sharp, K.A., Rastogi, T. & Axelsen, P.H. Intrinsic Linear Heterogeneity of Amyloid β Protein Fibrils Revealed by Higher Resolution Mass-per-length Determinations *. Journal of Biological Chemistry 285, 41843–41851 (2010).

86. Sen, A. et al. Mass analysis by scanning transmission electron microscopy and electron diffraction validate predictions of stacked beta-solenoid model of HET-s prion fibrils. J Biol Chem 282, 5545–50 (2007).

87. Tang, G. et al. EMAN2: An extensible image processing suite for electron microscopy. Journal of Structural Biology 157, 38–46 (2007).

88. Iadanza, M.G., Jackson, M.P., Radford, S.E. & Ranson, N.A. MpUL-multi: Software for Calculation of Amyloid Fibril Mass per Unit Length from TB-TEM Images. Scientific Reports 6, 21078 (2016).

89. Phillips, J.C. et al. Scalable molecular dynamics on CPU and GPU architectures with NAMD. The Journal of Chemical Physics 153, 044130 (2020).

90. Huang, J. et al. CHARMM36m: an improved force field for folded and intrinsically disordered proteins. Nature Methods 14, 71–73 (2017).

91. Feller, S.E., Zhang, Y., Pastor, R.W. & Brooks, B.R. Constant pressure molecular dynamics simulation: The Langevin piston method. The Journal of Chemical Physics 103, 4613–4621 (1995).

92. Martyna, G.J., Tobias, D.J. & Klein, M.L. Constant pressure molecular dynamics algorithms. The Journal of Chemical Physics 101, 4177–4189 (1994).

93. Darden, T., York, D. & Pedersen, L. Particle mesh Ewald: An N⋅log(N) method for Ewald sums in large systems. The Journal of Chemical Physics 98, 10089–10092 (1993).

