## Supplementary notes and extended data figures for "Structure of alpha-synuclein fibrils derived from human Lewy body dementia tissue"

#### Chemical Shift Assignments

Chemical shift assignments were performed for the LBD tissue-derived Asyn fibril using two independently produced samples — one with uniform [$^{13}C$, $^{15}N$] labeling (uCN) and a second with uniform [$^{13}C$, $^{2}H$, $^{15}N$] labeling (uCDN) followed by back-exchange in 99.8% $D_2O$ after fibril amplification done in 100% $H_2O$. Resonance assignments for the uCN sample were determined using 2D $^{13}C$-$^{13}C$, 3D $^{15}N$-$^{13}C\alpha$-$^{13}CX$, 3D $^{15}N$-$^{13}C'$-$^{13}CX$, and 3D $^{13}C\alpha$-$^{15}N$-$^{13}C'$ data sets following standard procedures[1,2]. The complete list of data sets used for performing assignments using $^{13}C$-detection is presented in Supplementary Table 5 and $^{13}C$-$^{15}N$ resonance assignments have been submitted to the BioMagRes Bank (BMRB accession code 51678). The 2D $^{13}C$-$^{13}C$ spectrum serves as a conformational fingerprint of the fibril and provides some initial insights into the structure (Fig. 3a). Globally, the resolved peaks display decoupled line widths of < 0.4 ppm in the direct $^{13}C$ dimension, similar to those observed for in vitro Asyn fibrils and indicative of a highly ordered core[3,4]. Particular residue types including most Thr, Val, and Gly residues, as well as select spin systems from other residues such as L38, N65, Q79, S87, I88, and F94 are particularly well resolved in the 2D. The Lys and Glu regions, which account for 20% of the primary sequence between residues 30 and 100, are less well resolved than in the in vitro forms, indicating disorder or partial mobility for some of these residues. In contrast to the in vitro form, the Ala regions are less well resolved in the 2D $^{13}C$-$^{13}C$, but the signals have comparable intensity to the resolved regions, indicating a high degree of order consistent with beta-strand conformations. Interestingly, some conformational disorder is evident for S87 and I88. The 2D $^{15}N$-$^{13}C'$ and $^{15}N$-$^{13}C\alpha$ spectra also serve as structural fingerprints for the fibril, with a focus on the backbone atoms (Fig. 1b-c). Critically, these highlight the benefit of adding a $^{15}N$ dimension to disambiguate shifts, particularly for the Ala and Gly regions, which comprise 30% of the primary sequence between residues 30 and 100, as well as for key core residues like V71, V74, T75 and V77. Overall, there appears to be a predominant defined conformation that exhibits some localized heterogeneity.

Resonance assignments for the uCDN sample were determined using 3D $^{1}H$-detection experiments including $^{13}C\alpha$-$^{15}N$- $^{1}H$, $^{13}C'$-($C\alpha$)-$^{15}N$- $^{1}H$, $^{13}C'$-$^{15}N$- $^{1}H$, and $^{13}C\alpha$-($C'$)-$^{15}N$- $^{1}H$ and performed as described[2,5]. The complete list of data sets used for performing assignments using $^{1}H$-detection is presented in Supplementary Table 6. The benefits to using $^{1}H$-detection are: (1) the sensitivity is approximately four-fold greater than for $^{13}C$-detection, (2) the amide $^{1}H$ chemical shift can be used in conjunction with backbone $^{13}C$ and $^{15}N$ shifts for predicting phi and psi dihedral angles, (3) in the presence of $D_2O$ as the hydration liquid, signals only arise from $^{1}Hs$ participating in hydrogen bonding, (4) it provides an independent method for determining the backbone chemical shift assignments that is complementary to the $^{13}C$-detection methods. In addition, $^{13}C$-detection can still be utilized for performing a 2D $^{13}C$-$^{13}C$ fingerprint analysis (Extended Data Fig. 2a-b). As denoted by the labels, the peak patterns are identical to those of the uCN sample, demonstrating fidelity of the amplification protocol. This similarity is also evident from the 2D $^{15}N$-$^{13}C'$ and $^{15}N$-$^{13}Ca$ projections from the 3D $^{13}Ca$-$^{15}N$- $^{1}H$ and $^{13}C'$-$^{15}N$- $^{1}H$ experiments (Fig, 1c-d). In addition to confirming the backbone $^{13}C$ and $^{15}N$ assignments determined for the uCN sample, additional assignments were made for residues K43, Q62, E83, G84, A85, and G86, which gave better resolved data in the $^{1}H$-detected experiments than in the $^{13}C$-detected 3Ds. All $^{13}C$ and $^{15}N$ atoms with unambiguous assignments from either or both samples are summarized as filled black circles on the primary sequence (Extended Data Fig. 3), with the corresponding chemical shift assignments in Supplementary Table S7.

Backbone phi and psi dihedral angles were predicted using assignments for $^{15}$N, $^{13}$Cα, $^{13}$C', $^{13}$Cβ, and $^{1}H^{N}$ using TALOS-N[6], and are consistent with a primarily beta-strand secondary structure (Fig. 3d). Relative peak intensities in a CP experiment provide a qualitative indicator of the relative rigidity for the different regions. This is plotted for the 3D $^{13}$Cα-$^{15}$N- $^{1}$H data (Fig. 2d), and in conjunction with the TALOS-N results, suggest a protomer structure composed of five beta-strands. Since the uCDN sample was back-exchanged in $D_2O$ after seed amplification, the weaker peaks in the $^{1}$H-detected data suggests residues that could be in loops or turns that are otherwise rigid, but not involved in strong backbone hydrogen bonding and may exchange slowly with the $D_2O$ hydrating liquid. Using the same logic, the residues that show strong peaks in the $^{1}$H-detected data must be participating in backbone hydrogen bonding between beta-strands.

**Distance restraints from solid-state NMR**

The next step towards determining the structural fold of the LBD Asyn fibril was to detect medium- ($2 \leq |i-j| \leq 4$) and long-range ($|i-j| \geq 5$) distance correlations to be used as restraints in a structure calculation. We performed measurements of distances using pulse sequences that minimize the effects of dipolar truncation[8] such as PAR[9]. This sequence takes advantage of a proton-drive second-order polarization transfer mechanism that is resilient to dipolar truncation and has been demonstrated to generate high-quality distance restraints for structure determinations of several proteins including a uniform [$^{13}$C, $^{15}$N] labeled amyloid-beta fibril[10]. Here, PAR mixing was incorporated into a 2D $^{13}$C-$^{13}$C pulse sequence and used to detect medium- and long-range correlations in the uCN LBD fibril sample (Extended Data Fig. 4). In comparison to the spectrum in Fig. 3, which was collected with 75 ms DARR mixing[11], the 12 ms PAR spectrum displays significantly more intensity arising from the medium and long range correlations, which encode the highest value structural information. Even the 2D spectrum exhibits a number of well resolved strips that have an unambiguous assignment in one dimension, including I88CD1, I88CG2, V70CB, G73CA, L38CA, S87CB, K43C, V66C, and Q79CD. Additionally, a substantial set of peaks have only two possible assignments in one dimension, including L38CD1 or V77CG2, L38CG or I88CG1, V74CB or Q79CB, N65CB or Lys CE, L38CB or G67CA, N65CA or H50CA, and G41C or G93C. Many additional correlations were observed but could not be assigned from the 2D spectrum alone.

To disambiguate some of the other regions of spectral overlap, PAR mixing was incorporated into 3D pulse sequences (Extended Data Fig. 5). We employed a modification of the 3D $^{13}$C-$^{13}$C-$^{13}$C pulse sequence from Hong and co-workers,[12] utilizing RFDR[13,14] for the first mixing period to maximize the intensity of one-bond correlations and then PAR transfer during the second mixing period to detect long range correlations. This combination of mixing elements gave especially strong and well resolved signals for Gly, Ala, Thr, Val, Asn and Gln residues. We also employed a related 3D $^{15}$N-$^{13}$C-$^{13}$C pulse sequence[15] utilizing constant-time evolution[16] in the $^{15}$N dimension to provide resolution enhancement, followed by SPECIFIC CP[17] to transfer magnetization selectively to either the Cα or C' spectral regions prior to PAR transfer between $^{13}$Cs. Both sequences utilized non-uniform sampling (NUS) and SMILE processing of the two indirect dimensions[18-20], reducing experimental times and enhancing the resolution and sensitivity, while allowing broadband sampling of all three $^{13}C$ dimensions.

Several long-range correlations were identified in the 2D $^{13}C$-$^{13}C$ experiment by exploiting the unambiguously assigned strips in the indirect dimension (Extended Data Fig. 6a). Substantial portions of the spectra could be assigned uniquely as a validation fo the data quality. For example, we assigned correlations from L38CA to A78CA/CB; V70CB to N65CA/CB; and G73CA to F94CD. Additionally, T92CB and T72CB are clearly resolved in the 3D $^{13}C$-$^{13}C$-$^{13}C$ and independently exhibit distinct cross peaks with F94CD (Extended Data Fig. 6b). The 2D also reveals cross peaks from S87CB to either A78CA or A91CA, as well as multiple cross peaks between the I88 methyl groups and the Q79 and V77 spin systems. Additional correlations from the 3D $^{13}C$-$^{13}C$-$^{13}C$ include cross peaks between the G41CA and A69CA/CB and V70CB; N65CB and V70CA/CB/CG; G68CA to S42CA/CB and G41C; T75CB to T92CB; and T92CB to V74C'/CA/CB and G73C'. Finally, the 3D $^{15}N$-$^{13}C\alpha$-$^{13}CX$ resolves additional cross peaks between S42CA and A69CA/CB and G68C'; A69CA and S42CA/CB and G41C'; V74CA and T92CB; and A78CA to L38, G36CA, and I88CG2.

In addition to the distance restraints generated using the uCN sample, a third sample was prepared using isotopic dilution; Asyn monomer prepared with a [2-$^{13}C$]-glycerol, uniform-$^{15}N$ labeling pattern was diluted approximately four-fold in natural abundance monomer prior to fibril amplification (diluted). The advantage of this sample was that long-range cross peaks in a 2D $^{13}C$-$^{13}C$ correlation experiment arise primarily from *intra*molecular interactions, since those arising from *inter*molecular interactions are expected to have four times lower peak intensities. Unambiguous long-range correlations such as between Q79CD and I88CD1 were used as important tertiary structure restraints (Extended Data Fig. 7).

Finally, a $^{15}N$-dephased, $^{13}C$-detected frequency-selective REDOR (FS-REDOR) experiment[21] performed on the uCN sample was used to observe the presence of salt bridge(s) between the Lys amine NZs and the Glu carboxylate CDs (Extended Data Fig. 8). As an internal control, the selective pulse on the $^{13}C$ channel was set to encompass both the Glu CDs and the carbonyl peak. With 1.8 ms REDOR mixing, an effect is not observed for either the Glu CDs or the carbonyls. However, with 9.0 ms REDOR mixing, a 30% dephasing is observed for the Glu CD peak, but not for the carbonyls (red spectrum), confirming that at least one (and likely multiple) salt bridge(s) must be present in the fibril structure.

### SSNMR structural model demonstrates high stability in an unrestrained molecular dynamics simulation

We next proceeded to analyze the thermodynamic stability of the SSNMR structure using an unrestrained molecular dynamics (MD) simulation. While the results of this simulation do not directly validate the accuracy of the model, they demonstrate clearly that the fibril is remarkably stable (Extended Data Fig. 9a-b), and regions where NMR data is sparse correlate to regions displaying a high degree of disordering throughout the simulation. The fibril core (residues 35 to 44 and 65 to 99) show an RMSD of ~3.6 Å and an average over the whole structure of ~5.75 Å, where the higher RMSD is contributed by the disorder of the N- and C- termni and the disordered residues 45 to 64. The RMSF measured for each residue was less than 1 Å for the fibril core

(Extended Data Fig. 9c) relative to the final NMR structure. After 200 ns of production MD, the overall fold remains qualitatively unchanged, however the fibril does inherit a twist (Extended Data Fig. 9d-e). The RMSD value for individual protomers relative to the SSNMR model stabilizes at about 2.5. Å, in close agreement with the uncertainty in the SSNMR bundle The RMSD calculated for the full fibril stabilizes at about 3.25 Å, but the inflation is primarily due to the twist. Additionally, we identified ions in the in the simulation coordinated to specific regions within the simulation. Extended Data Fig. 9f. shows a projection along the long axis of the fibril where the ion distribution of sodium coordinate to the C-terminus and to E83. Extended Data Fig. 9g shows chloride ions coordinating predominantly to residues 32 to 45, where two KTK repeats are present within the structure, with the lysine sidechains coordinating to the chloride ion mass. This is interesting as in the cryo-EM structure of LBD fibrils display a non-proteinaceous density at this same site.

**Supplementary Table 1**: **Concentration of amplified fibrils produced with natural abundance monomer, as determined by the [$^3$H]-BF2846 radioligand binding assay.**

| | After 6$^{th}$ cycle | |
|---|---|---|
| **Case** | **Concentration (µM)** | **Fold amplification** |
| LBD1 | 92.6 | 17.1 |
| LBD2 | 94.2 | 17.4 |
| LBD3 | 87.3 | 16.1 |
| LBD4 | 189.3 | 34.9 |
| C1 | 4.0 | |
| C2 | 10.1 | |
| C3 | 2.2 | |

**Supplementary Table 2**: **Concentration of amplified fibrils produced with isotopically labeled Asyn monomer (uCN-Asyn), as determined by the [$^3$H]-BF2846 radioligand binding assay.**

| | After 4$^{th}$ cycle | | After 6$^{th}$ cycle | |
|---|---|---|---|---|
| **Case** | **Concentration (µM)** | **Fold amplification** | **Concentration (µM)** | **Fold amplification** |
| LBD1 | 39.83 | 22.8 | 42.24 | 11.5 |
| LBD2 | 19.65 | 11.2 | | |
| C1 | 2.88 | | 3.15 | |
| C2 | 0.62 | | 4.19 | |

**Supplementary Table 3**: **Concentration of amplified fibrils produced with isotopically labeled Asyn monomer (CDN-Asyn), as determined by the [$^{3}$H]-BF2846 radioligand binding assay.**

| | After 4$^{th}$ cycle | | After 6$^{th}$ cycle | |
|---|---|---|---|---|
| **Case** | **Concentration (µM)** | **Fold amplification** | **Concentration (µM)** | **Fold amplification** |
| LBD1 | 20.19 | 30.9 | 26.92 | 17.8 |
| LBD2 | 12.50 | 19.1 | | |
| C1 | 0.99 | | 0.26 | |
| C2 | 0.31 | | 2.76 | |

**Supplementary Table 4**: **Concentration of amplified fibrils produced with uCN and CDN isotopically labeled monomer, as determined by the micro-BCA assay.**

| | | After 6$^{th}$ cycle | |
|---|---|---|---|
| **Case** | **Asyn Monomer** | **Fibril concentration (µM)** | **Total yield (µg)** |
| LBD1 | uCN-Asyn | 43.4 (average of 10 tubes) | 5035 |
| C1 | uCN-Asyn | 6.4 | 73.4 |
| C2 | uCN-Asyn | 5 | 58 |
| LBD1 | CDN-Asyn | 33.4 (average of 10 tubes) | 3816 |
| C1 | CDN-Asyn | 2.2 | 25.1 |
| C2 | CDN-Asyn | 2.3 | 26.5 |

**Supplementary Table 5: Human postmortem brain tissue cases used for the study.** Grey matter from caudate brain region was selected for amplification.

| Case | Gender | Age at Death (years) | Post-Mortem interval (hours) | Primary Diagnosis |
|---|---|---|---|---|
| LBD1 | male | 71 | 14 | LBD |
| LBD2 | male | 76 | 4 | LBD |
| LBD3 | male | 76 | n/a | LBD |
| LBD4 | male | 78 | 14 | LBD |
| LBD5 | female | 82 | 3 | LBD |
| MSA1 | male | 58 | 22 | MSA |
| MSA2 | male | 58 | 12 | MSA |
| MSA3 | male | 65 | 7 | MSA |
| MSA4 | female | 59 | 5 | MSA |
| C1 | male | 85 | 30 | disease-control |
| C2 | female | 78 | 28 | disease-control |
| C3 | male | 73 | 22 | disease-control |

### Supplementary Table 6: Solid-state NMR experiments

| Sample | Experiment | Magnet | MAS Rate (kHz) | Mixing Time (ms) | NUS sampling density (%) | Exp. Time (hr) |
|---|---|---|---|---|---|---|
| **uCN** | 2D $^{13}C$-$^{13}C$ with DARR mixing | 17.6 T (WB) | 16.667 | 75, 300, 500 | N/A | 2.8, 76.8, 92.0 |
| **uCN** | 2D $^{13}C$-$^{13}C$ with SPC7 mixing | 17.6 T (WB) | 16.667 | 0.96 | N/A | 20.9 |
| **uCN** | 3D $^{15}N$-$^{13}CA$-$^{13}CX$ with DARR mixing | 17.6 T (WB) | 16.667 | 75 | 25 | 37.8 |
| **uCN** | 3D $^{15}N$-$^{13}CO$-$^{13}CX$ with DARR mixing | 17.6 T (WB) | 16.667 | 75 | 12.5 | 47.3 |
| **uCN** | 3D $^{13}CA$-$^{15}N$- $^{13}CO$ | 17.6 T (WB) | 16.667 | N/A | N/A | 90.4 |
| **uCN** | 2D $^{13}C$-$^{13}C$ with DARR mixing | 11.7 T (WB) | 22.222 | 50, 200, 500 | N/A | 5.4, 39.9, 130.6 |
| **uCN** | 2D $^{15}N$-$^{13}CA$ | 11.7 T (WB) | 22.222 | N/A | N/A | 4.4 |
| **uCN** | 2D $^{15}N$-$^{13}CO$ | 11.7 T (WB) | 22.222 | N/A | N/A | 0.9 |
| **uCN** | 3D $^{15}N$-$^{13}CA$-$^{13}CX$ with DARR mixing | 11.7 T (WB) | 22.222 | 200 | 12.5 | 363 |
| **uCN** | 3D $^{15}N$-$^{13}CO$-$^{13}CX$ with DARR mixing | 11.7 T (WB) | 22.222 | 200 | 12.5 | 152 |
| **uCN** | 3D $^{15}N$-$^{13}CO$-$^{13}CA$ with SPC5 mixing | 11.7 T (WB) | 22.222 | 1.26 | 12.5 | 61.4 |
| **uCN** | 3D $^{13}CA$-$^{15}N$- $^{13}CO$ | 11.7 T (WB) | 22.222 | N/A | 12.5 | 41.0 |
| **uCN** | 2D $^{13}C$-$^{13}C$ with PAR mixing | 17.6 T (WB) | 17.007 | 12.9 | N/A | 56.8 |
| **uCN** | 3D $^{13}C$-$^{13}C$- $^{13}C$ with RFDR then PAR mixing | 17.6 T (WB) | 17.241 | 1.86 then 12.8 | 25 | 227.2 |
| **uCN** | 3D $^{15}N$-$^{13}CA$-$^{13}CX$ with PAR mixing | 11.7 T (WB) | 22.222 | 12.7 | 10 | 446 |
| **uCN** | 1D $^{13}C$ with fsREDOR mixing | 11.7 T (WB) | 11.111 | 1.8, 5.4, 9.0, 10.8 | N/A | 1.4 |
| **uDCN** | 3D $^{13}CA$-$^{15}N$-$^{1}H$ | 17.6 T (WB) | 33.333 | N/A | N/A | 44.1 |
| **uDCN** | 3D $^{13}CO$-$^{15}N$-$^{1}H$ | 17.6 T (WB) | 33.333 | N/A | N/A | 66.5 |
| **uDCN** | 3D $^{13}CA$-{$^{13}CO$}- $^{15}N$-$^{1}H$ with RFDR mixing | 17.6 T (WB) | 33.333 | 1.92 | 12.5 | 37.2 |
| **uDCN** | 3D $^{13}CO$-{$^{13}CA$}- $^{15}N$-$^{1}H$ with RFDR mixing | 17.6 T (WB) | 33.333 | 1.92 | 12.5 | 57.8 |
| **uDCN** | 3D $^{13}CA$-$^{15}N$-{$^{1}H$}-$^{1}H$ with RFDR mixing | 17.6 T (WB) | 33.333 | 7.68 | N/A | 41.5 |
| **uDCN** | 3D $^{13}CA$-{$^{1}H$ -$^{1}H$}-$^{15}N$-$^{1}H$ with RFDR mixing | 17.6 T (WB) | 33.333 | 7.68 | N/A | 59.1 |
| **uDCN** | 3D $^{15}N$-{$^{1}H$ -$^{1}H$}-$^{15}N$-$^{1}H$ with RFDR mixing | 17.6 T (WB) | 33.333 | 7.68 | N/A | 97.8 |
| **uDCN** | 4D $^{1}H$-$^{15}N$-{$^{1}H$ -$^{1}H$}-$^{15}N$-$^{1}H$ with RFDR mixing | 17.6 T (WB) | 33.333 | 7.68 | 3 | 85.2 |
| **uDCN** | 2D $^{13}CA$-$^{13}CO$ with CTUC-COSY mixing | 17.6 T (WB) | 33.333 | 9.6 | N/A | 7.1 |
| **uDCN** | 2D $^{13}Cali$-$^{13}Cali$ with fpRFDR mixing | 11.7 T (WB) | 33.333 | 11.78 | N/A | 43.5 |
| **diluted** | 2D $^{13}C$-$^{13}C$ with DARR mixing | 11.7 T (WB) | 11.111 | 500 | N/A | 277.9 |
| **diluted** | 2D $^{15}N$-$^{13}CA$ | 11.7 T (WB) | 11.111 | N/A | N/A | 16.3 |
| **diluted** | 2D $^{15}N$-$^{13}CO$ | 11.7 T (WB) | 11.111 | N/A | N/A | 18.7 |
| | | | | | **Total Time** | 2750 |

**Supplementary Table 7: $^{13}$C, $^{15}$N, and $^{1}$H$^{N}$ chemical shift assignments for LBD derived a-synuclein fibrils**

| Residue | $^{1}$H$^{N}$ | $^{15}$N | $^{13}$C’ | $^{13}$CA | $^{13}$CB | $^{13}$CG | $^{13}$CD | $^{13}$CE | $^{13}$CZ | $^{15}$ND/$^{15}$NE/$^{15}$NZ |
|---|---|---|---|---|---|---|---|---|---|---|
| -- | -- | -- | -- | -- | -- | -- | -- | -- | -- | -- |
| K34 | 7.46 | 125.5 | 174.5 | 56.1 | - | - | - | - | - | - |
| E35 | 8.76 | 121.9 | 176.3 | 53.4 | 35.7 | 35.6 | 183.4 | | | |
| G36 | 8.75 | 117.5 | 173.6 | 48.4 | | | | | | |
| V37 | 7.67 | 118.6 | 172.4 | 60.0 | 35.7 | 21.9/24.5 | | | | |
| L38 | 8.88 | 127.8 | 173.8 | 52.6 | 47.3 | 27.5 | 25.3/29.0 | | | |
| Y39 | 8.39 | 131.3 | 173.9 | 56.3 | 43.4 | 128.0 | 133.0 | 117.5 | 157.3 | |
| V40 | 8.76 | 127.1 | 173.8 | 60.0 | 34.4 | 19.9/22.1 | | | | |
| G41 | 9.08 | 111.6 | 170.2 | 44.7 | | | | | | |
| S42 | 8.94 | 116.9 | 175.4 | 56.5 | 68.7 | | | | | |
| K43 | 7.96 | 123.4 | 176.6 | 56.6 | - | - | - | - | - | - |
| -- | -- | -- | -- | -- | -- | -- | -- | -- | -- | -- |
| V49 | - | - | - | - | - | | | | | |
| H50 | - | - | 175.8 | 51.7 | 31.5 | 125.1 | 129.4 | 140.3 | | -/- |
| -- | -- | -- | -- | -- | -- | -- | -- | -- | -- | -- |
| E61 | - | - | 173.1 | - | - | - | - | | | |
| Q62 | 8.15 | 122.6 | 174.3 | 54.5 | - | - | - | | | - |
| V63 | 8.96 | 124.6 | 173.9 | 61.2 | 35.5 | 21.1/- | | | | |
| T64 | 9.32 | 124.8 | 172.3 | 60.8 | 68.5 | 23.4/23.4 | | | | |
| N65 | 9.61 | 126.5 | 173.0 | 51.9 | 42.0 | 174.4 | | | | 113.9 |
| V66 | 9.32 | 125.0 | 177.0 | 59.9 | 34.4 | 20.9/23.0 | | | | |
| G67 | 8.83 | 112.6 | 171.9 | 47.4 | | | | | | |
| G68 | 6.07 | 100.7 | 172.1 | 44.4 | | | | | | |
| A69 | 8.74 | 121.8 | 175.0 | 50.0 | 22.1 | | | | | |
| V70 | 8.95 | 124.5 | 175.8 | 61.7 | 32.7 | 19.7/21.2 | | | | |
| V71 | 9.18 | 131.0 | 173.9 | 61.3 | 34.7 | 19.7/22.1 | | | | |
| T72 | 9.28 | 124.7 | 174.6 | 60.1 | 71.4 | 22.5 | | | | |
| G73 | 8.96 | 117.8 | 172.1 | 49.0 | | | | | | |
| V74 | 7.73 | 117.6 | 173.8 | 59.4 | 37.4 | 20.3/23.3 | | | | |
| T75 | 8.86 | 123.4 | 171.8 | 61.7 | 68.8 | 22.9 | | | | |
| A76* | 9.55 | 131.2 | 174.2 | 49.7 | 21.7 | | | | | |
| V77* | 8.76 | 122.4 | 172.2 | 60.6 | 36.7 | 20.6/25.1 | | | | |
| A78 | 9.17 | 130.3 | 175.7 | 50.7 | 23.1 | | | | | |
| Q79 | 8.80 | 122.6 | 174.1 | 54.6 | 37.6 | 34.7 | 179.3 | | | 112.4 |
| K80 | 8.97 | 126.5 | 173.9 | 54.7 | - | - | - | - | - | - |
| T81 | 8.53 | 124.7 | 172.8 | 61.9 | 69.9 | 21.1 | | | | |
| V82 | 9.42 | 127.0 | 175.4 | 60.1 | - | -/- | | | | |
| E83 | 8.93 | 118.3 | 175.9 | 55.3 | | | | | | |
| G84 | 8.96 | 124.4 | 175.0 | 49.8 | | | | | | |
| A85 | 9.13 | 124.7 | 174.5 | 49.9 | - | | | | | |
| G86 | 8.86 | 110.3 | 172.5 | 44.2 | | | | | | |
| S87* | 9.46 | 119.7 | 174.4 | 56.4 | 67.4 | | | | | |
| I88* | 8.96 | 128.2 | 173.2 | 60.1 | 44.4 | 27.8/17.6 | 16.9 | | | |
| A89 | 8.97 | 129.1 | 175.2 | 50.3 | 22.5 | | | | | |
| A90 | 9.31 | 126.3 | 174.4 | 49.7 | 22.2 | | | | | |
| A91 | 8.73 | 123.7 | 175.4 | 50.3 | 23.2 | | | | | |
| T92 | 8.79 | 117.3 | 173.3 | 61.2 | 70.8 | 20.5 | | | | |
| G93 | - | 116.8 | 170.3 | 44.9 | | | | | | |
| F94 | 9.16 | 129.0 | 173.1 | 56.9 | 43.5 | 137.4 | 131.2 | 129.3 | 140.3 | |
| V95 | 9.00 | 128.5 | 173.0 | 60.9 | 35.6 | -/- | | | | |
| K96 | 8.99 | 129.0 | 175.1 | 54.5 | - | - | - | - | - | - |
| -- | -- | -- | -- | -- | -- | -- | -- | -- | -- | -- |

(*) Two sets of shifts observed for at least one atom in the residue. Primary shift is reported here.

**Supplementary Table 8: Summary of simulated annealing protocols in the PASD algorithm**

| | pass 2 | pass 3,4,5 |
|---|---|---|
| *High temperature phase I* | | |
| Duration (ps) | 2 | 5 |
| $k_{\mathrm{lin}}$ (kcal/mol/Å) | 1 | 0 |
| $k_{\mathrm{quad}}$ (kcal/mol/Å$^2$) | 0 | 3 |
| $k_{\mathrm{repul}}$ (kcal/mol/Å) | 5 | 5 |
| $\Delta r_c$ (Å) | ∞ | ∞ |
| $w_p$ | 1 | 1 |
| Number of NOE re-evaluations | 64 | 64 |
| Number of MC steps | 25 | 25 |
| $k_{\mathrm{nb}}$ (kcal/mol/Å$^4$) | 0 | 0.004 |
| $s_{\mathrm{nb}}$ | | 1.2 |
| Nonbonded interactions | None | Cα-Cα only |
| $k_{R\mathrm{gyr}}$ (kcal/mol/Å$^3$) | 0.001 | 0.001 |
| $k_{\mathrm{dihed}}$ (kcal/mol/radians$^2$) | 200 | 10 |
| $k_{\mathrm{tDB}}$ (kcal/mol) | 0.002 | 0.002 |
| *High temperature phase II* | | |
| Duration (ps) | 6 | |
| $k_{\mathrm{lin}}$ (kcal/mol/Å) | 1 | |
| $k_{\mathrm{quad}}$ (kcal/mol/Å$^2$) | 0 | |
| $k_{\mathrm{repul}}$ (kcal/mol/Å) | 5 | |
| $\Delta r_c$ (Å) | 10 | |
| $w_p$ | 0.5 | |
| Number of NOE re-evaluations | 64 | |
| Number of MC steps | 25 | |
| $k_{\mathrm{nb}}$ (kcal/mol/Å$^4$) | 0 | |
| $s_{\mathrm{nb}}$ | | |
| Nonbonded interactions | None | |
| $k_{R\mathrm{gyr}}$ (kcal/mol/Å$^3$) | 0.001 | |
| $k_{\mathrm{dihed}}$ (kcal/mol/radians$^2$) | 200 | |
| $k_{\mathrm{tDB}}$ (kcal/mol) | 0.002 | |
| *Cooling phase* | | |
| Duration (ps) | 250 | 250 |
| $k_{\mathrm{lin}}$ (kcal/mol/Å) | 1 → 30 | 0 |
| $k_{\mathrm{quad}}$ (kcal/mol/Å$^2$) | 0 | 3 → 30 |
| $k_{\mathrm{repul}}$ (kcal/mol/Å) | 5 | 5 |
| $\Delta r_c$ (Å) | 10 → 2 | 2 → 0.7 |
| $w_p$ | 0.5 → 0 | 0.5 → 0 |
| Number of NOE re-evaluations | 128 | 128 |
| Number of MC steps | 100 | 100 |
| $k_{\mathrm{nb}}$ (kcal/mol/Å$^4$) | 0.004 → 1 | 0.004 → 1 |
| $s_{\mathrm{nb}}$ | 2.0 | 2.0 |
| Nonbonded interactions | All atoms | All atoms |
| $k_{R\mathrm{gyr}}$ (kcal/mol/Å$^3$) | 0.001 → 1.25 | 0.001 → 1.25 |
| $k_{\mathrm{dihed}}$ (kcal/mol/radians$^2$) | 200 | 10 → 300 |
| $k_{\mathrm{tDB}}$ (kcal/mol) | 0.002 → 2 | 0.002 → 2 |

### References

1. Comellas, G. & Rienstra, C.M. Protein structure determination by magic-angle spinning solid-state NMR, and insights into the formation, structure, and stability of amyloid fibrils. *Annu Rev Biophys* **42**, 515-36 (2013).
2. Higman, V.A. Solid-state MAS NMR resonance assignment methods for proteins. *Prog Nucl Magn Reson Spectrosc* **106-107**, 37-65 (2018).
3. Barclay, A.M., Dhavale, D.D., Courtney, J.M., Kotzbauer, P.T. & Rienstra, C.M. Resonance assignments of an α-synuclein fibril prepared in Tris buffer at moderate ionic strength. *Biomol NMR Assign* **12**, 195-199 (2018).
4. Comellas, G., Lopez, J.J., Nieuwkoop, A.J., Lemkau, L.R. & Rienstra, C.M. Straightforward, effective calibration of SPINAL-64 decoupling results in the enhancement of sensitivity and resolution of biomolecular solid-state NMR. *Journal of Magnetic Resonance* **209**, 131-135 (2011).
5. Zhou, D.H. & Rienstra, C.M. High-performance solvent suppression for proton detected solid-state NMR. *J Magn Reson* **192**, 167-72 (2008).
6. Shen, Y. & Bax, A. Protein backbone and sidechain torsion angles predicted from NMR chemical shifts using artificial neural networks. *J Biomol NMR* **56**, 227-41 (2013).
7. Castellani, F. et al. Structure of a protein determined by solid-state magic-angle-spinning NMR spectroscopy. *Nature* **420**, 98-102 (2002).
8. Bayro, M.J. et al. Dipolar truncation in magic-angle spinning NMR recoupling experiments. *J Chem Phys* **130**, 114506 (2009).
9. De Paëpe, G., Lewandowski, J.R., Loquet, A., Böckmann, A. & Griffin, R.G. Proton assisted recoupling and protein structure determination. *J Chem Phys* **129**, 245101 (2008).
10. Colvin, M.T. et al. Atomic Resolution Structure of Monomorphic Aβ42 Amyloid Fibrils. *J Am Chem Soc* **138**, 9663-74 (2016).
11. Takegoshi, K., Nakamura, S. & Terao, T. 13C–1H dipolar-assisted rotational resonance in magic-angle spinning NMR. *Chemical Physics Letters* **344**, 631-637 (2001).
12. Li, S., Zhang, Y. & Hong, M. 3D (13)C-(13)C-(13)C correlation NMR for de novo distance determination of solid proteins and application to a human alpha-defensin. *J Magn Reson* **202**, 203-10 (2010).
13. Bennett, A.E., Griffin, R.G., Ok, J.H. & Vega, S. Chemical shift correlation spectroscopy in rotating solids: Radio frequency-driven dipolar recoupling and longitudinal exchange. *The Journal of Chemical Physics* **96**, 8624-8627 (1992).
14. Bennett, A.E. et al. Homonuclear radio frequency-driven recoupling in rotating solids. *The Journal of Chemical Physics* **108**, 9463-9479 (1998).
15. Rienstra, C.M., Hohwy, M., Hong, M. & Griffin, R.G. 2D and 3D 15N−13C−13C NMR Chemical Shift Correlation Spectroscopy of Solids: Assignment of MAS Spectra of Peptides. *Journal of the American Chemical Society* **122**, 10979-10990 (2000).
16. Vuister, G.W. & Bax, A. Resolution enhancement and spectral editing of uniformly 13C-enriched proteins by homonuclear broadband 13C decoupling. *Journal of Magnetic Resonance (1969)* **98**, 428-435 (1992).
17. Baldus, M., Petkova, A.T., Herzfeld, J. & Griffin, R.G. Cross polarization in the tilted frame: assignment and spectral simplification in heteronuclear spin systems. *Molecular Physics* **95**, 1197-1207 (1998).
18. Zambrello, M.A. et al. Nonuniform sampling in multidimensional NMR for improving spectral sensitivity. *Methods* **138-139**, 62-68 (2018).

19. Paramasivam, S. et al. Enhanced sensitivity by nonuniform sampling enables multidimensional MAS NMR spectroscopy of protein assemblies. *J Phys Chem B* **116**, 7416-27 (2012).
20. Hoch, J.C., Maciejewski, M.W., Mobli, M., Schuyler, A.D. & Stern, A.S. Nonuniform Sampling in Multidimensional NMR. in *eMagRes* (2012).
21. Jaroniec, C.P., Tounge, B.A., Herzfeld, J. & Griffin, R.G. Frequency Selective Heteronuclear Dipolar Recoupling in Rotating Solids: Accurate 13C−15N Distance Measurements in Uniformly 13C,15N-labeled Peptides. *Journal of the American Chemical Society* **123**, 3507-3519 (2001).

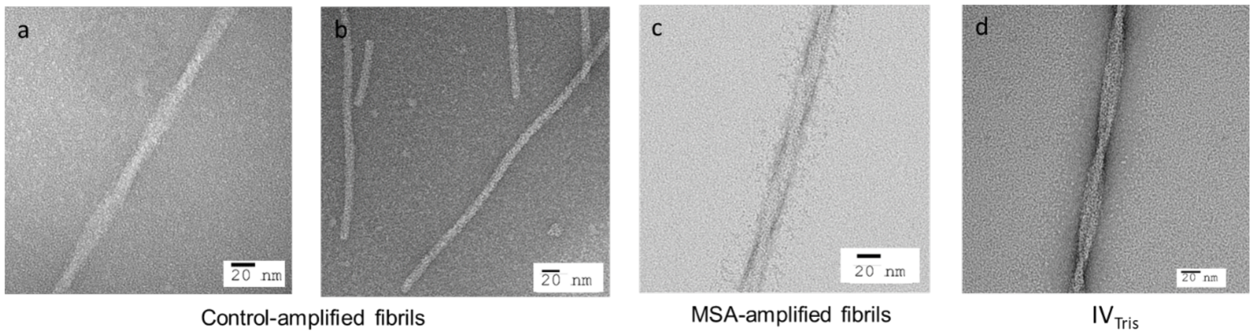

**Extended Data Fig. 1:** Negative stain TEM images of Asyn fibril preparations. Representative negative stain TEM images of control amplified fibrils (**a-b**), MSA (**c**) amplified fibrils, and in vitro fibrils preparared under nucleation conditions in Tris buffer ($IV_{Tris}$) (**d**).

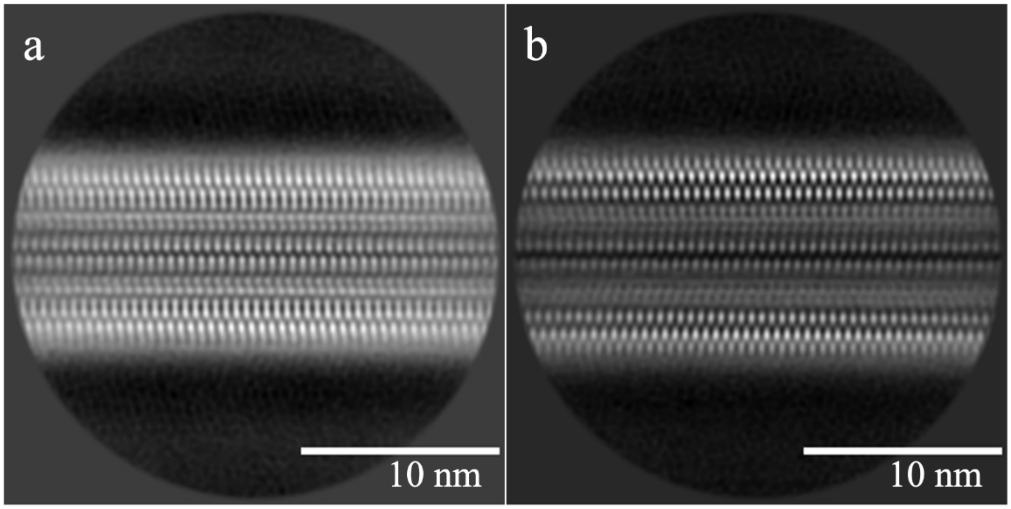

**Extended Data Fig. 2:** Cryo-EM 2D class averages for LBD amplified fibrils. **a-b,** Examples of 2D class averages obtained from single particle cryo-EM data. Both images show non-twisting fibrils that have features consistent with a two-protofilament structure and pseudo-$2_1$ helical screw symmetry.

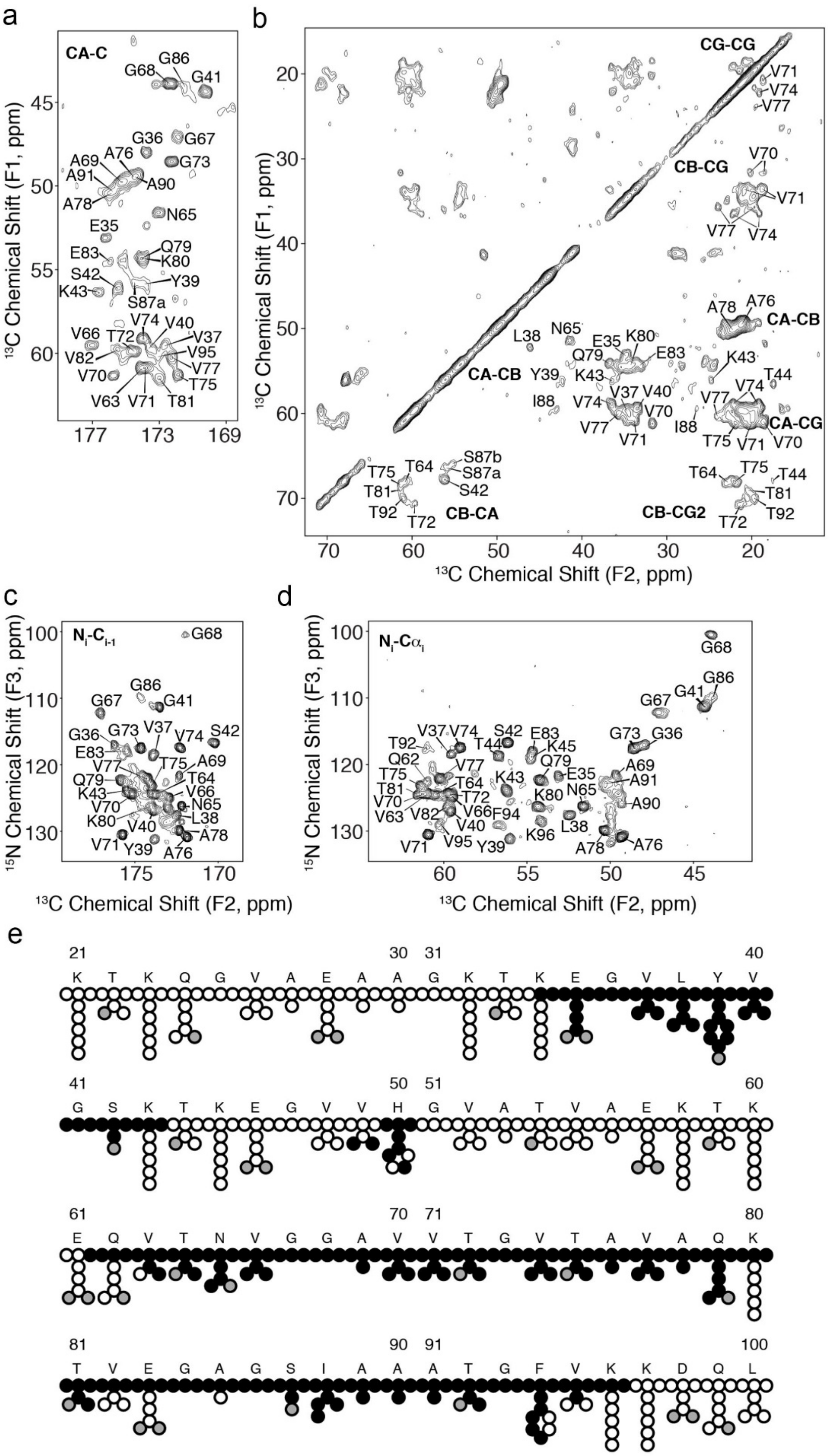

a
CA-C
G68
G86
G41
G36
G67
G73
A69
A76
A91
A90
A78
N65
E35
E83
Q79
K80
S42
Y39
K43
S87a
V74
V40
V66
T72
V37
V95
V82
V77
V70
T75
V63
V71
T81
$^{13}$C Chemical Shift (F1, ppm)
b
CG-CG
CB-CG
CA-CB
CB-CA
CB-CG2
CA-CG
L38
N65
E35
K80
Q79
E83
K43
Y39
V37
V40
V74
V70
I88
V77
V71
A78
A76
T44
V74
T75
S87b
S87a
S42
T64
T81
T92
T72
$^{13}$C Chemical Shift (F1, ppm)
$^{13}$C Chemical Shift (F2, ppm)
c
$N_i$-$C_{i-1}$
G68
G86
G41
G67
G36
G73
V37
V74
S42
E83
A69
V77
T75
Q79
T64
K43
V66
N65
V70
L38
K80
V40
A78
V71
Y39
A76
$^{15}$N Chemical Shift (F3, ppm)
$^{13}$C Chemical Shift (F2, ppm)
d
$N_i$-$C\alpha_i$
G68
G86
G41
G67
G73
G36
V37
V74
S42
E83
T92
T44
K45
Q62
V77
A69
K43
Q79
E35
A91
T75
T64
T81
T72
V70
K80
N65
A90
V63
V82
V66
F94
L38
V40
V71
V95
Y39
K96
A78
A76
$^{15}$N Chemical Shift (F3, ppm)
$^{13}$C Chemical Shift (F2, ppm)
e
21
K T K Q G V A E A A
30
31
G K T K E G V L Y V
40
41
G S K T K E G V V H
50
51
G V A T V A E K T K
60
61
E Q V T N V G G A V
70
71
V T G V T A V A Q K
80
81
T V E G A G S I A A
90
91
A T G F V K K D Q L
100

**Extended Data Fig. 3:** Two-dimensional SSNMR spectra collected on uniformly $^{2}H$,$^{13}C$,$^{15}N$-labeled LBD Asyn fibril. **a**, $^{13}C$-$^{13}C$ spectrum showing alpha to carbonyl correlations. Data was acquired at 750 MHz $^{1}H$ frequency with 33.333 kHz magic-angle spinning and sample temperature 10±5 ºC with 9 ms CTUC-COSY mixing. The indirect dimension was extended two-fold during processing using SMILE. **b**, $^{13}C$-$^{13}C$ spectrum showing aliphatic correlations. Data was acquired at 500 MHz $^{1}H$ frequency with 22.222 kHz magic angle spinning and a variable temperature set point of 0 ºC with 11.8 ms fpRFDR mixing. **c**, $^{15}N$-$^{13}C$ projection from a 3D CONH experiment showing backbone correlations between amide nitrogens and carbonyl carbons. **d**, $^{15}N$-$^{13}C$ projection from a 3D CANH spectrum showing backbone correlations between amide nitrogens and alpha carbons. Data in (**b**) and (**c**) were acquired at 750 MHz $^{1}H$ frequency with 33.333 kHz magic-angle spinning and sample temperature 10±5 ºC, using a 6 ms $^{15}N$-$^{13}C$ cross polarization. 3Ds were collected using non-uniform sampling followed by reconstruction using SMILE prior to Fourier transformation. **e**, Assigned atoms showing residues K21 to L100 out of 140 total amino acids in the Asyn primary sequence. Atoms with assignments are represented as filled black circles, filled gray circles are oxygen, and empty circles are unassigned or left empty due to uncertainty in the assignment.

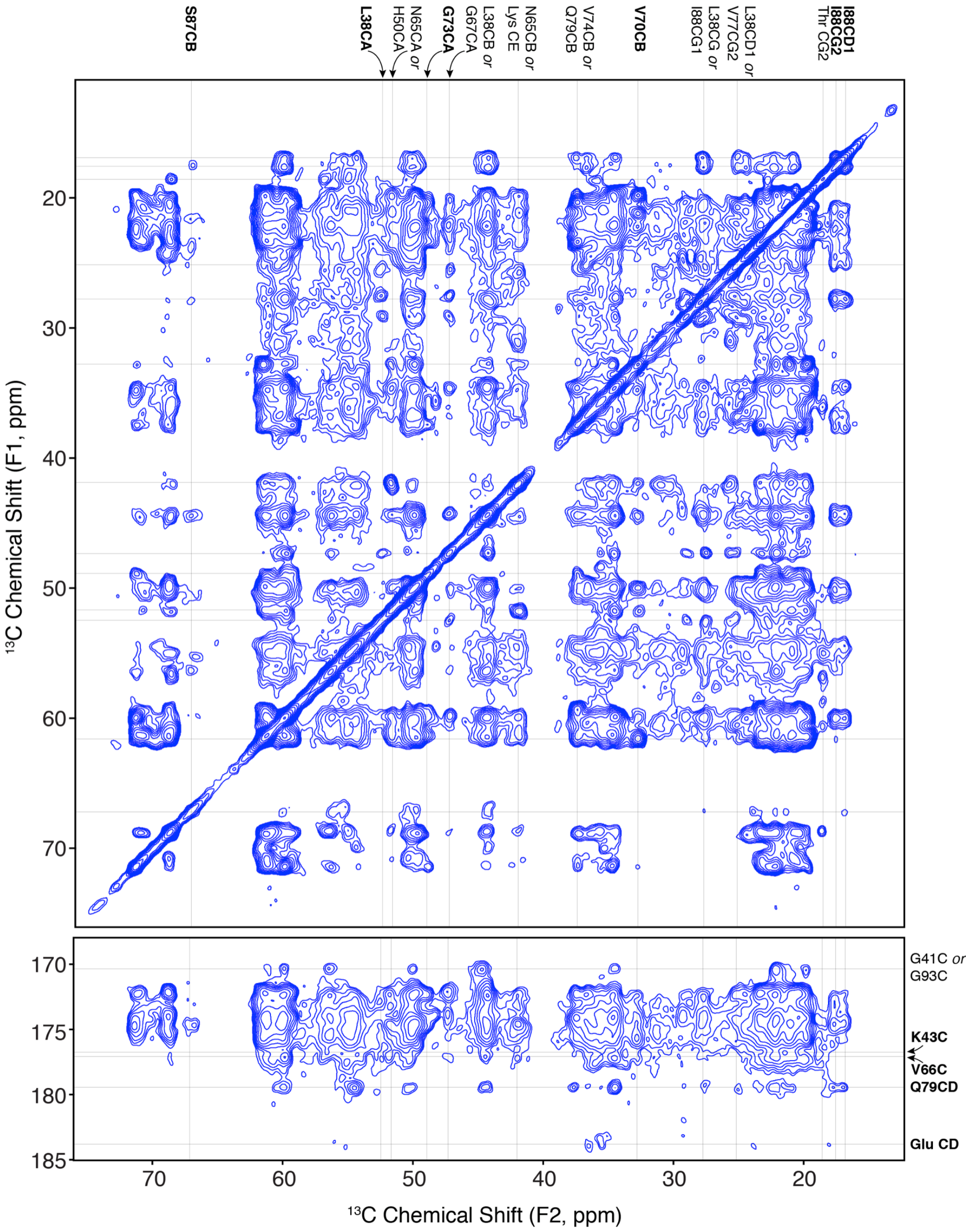

**Extended Data Fig. 4:** Two-dimensional $^{13}$C-$^{13}$C correlation spectrum acquired with PAR mixing on uniformly $^{13}$C,$^{15}$N-labeled LBD Asyn fibril. $^{13}$C-$^{13}$C correlation spectrum showing the aliphatic-aliphatic (top panel) and aliphatic-carbonyl (bottom panel) regions. Data was acquired at 750 MHz $^{1}$H frequency with 16.667 kHz magic-angle spinning with 12 ms PAR mixing. The first contour is cut at 6 times the root-mean-square noise. Horizontal and vertical lines are drawn to indicate all strips with one of the two assignments being either unambiguous (Bold atom labels) or two-fold ambiguous (regular text labels).

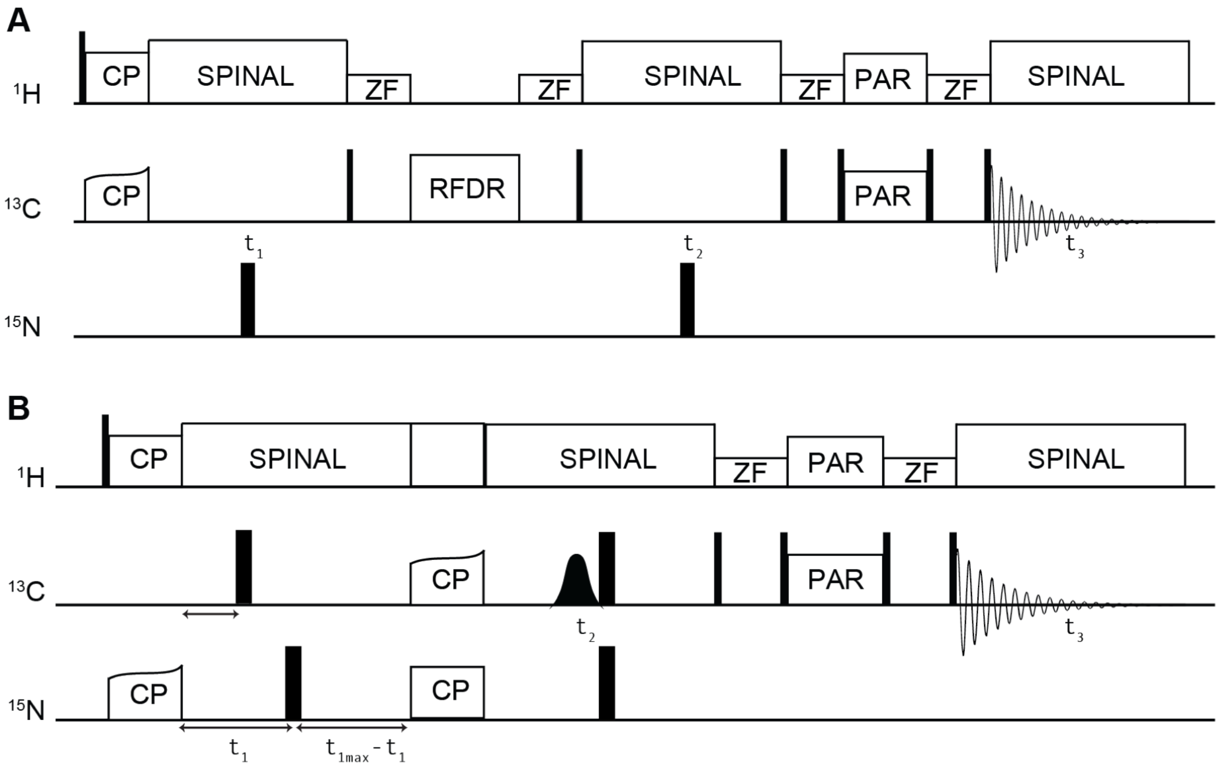

**Extended Data Fig. 5:** Pulse sequence diagrams for 3D experiments with PAR mixing. **a**, Pulse sequence used to collect broadband $^{13}C$-$^{13}C$-$^{13}C$ correlation spectrum. **b**, Pulse sequence used to collect $^{15}N$-$^{13}C\alpha$-$^{13}CX$ correlation spectrum utilizing constant-time evolution for $^{15}N$ and a J-decoupling soft pulse for $^{13}C\alpha$. Both experiments can be done using non-uniform sampling of the two indirect dimensions.

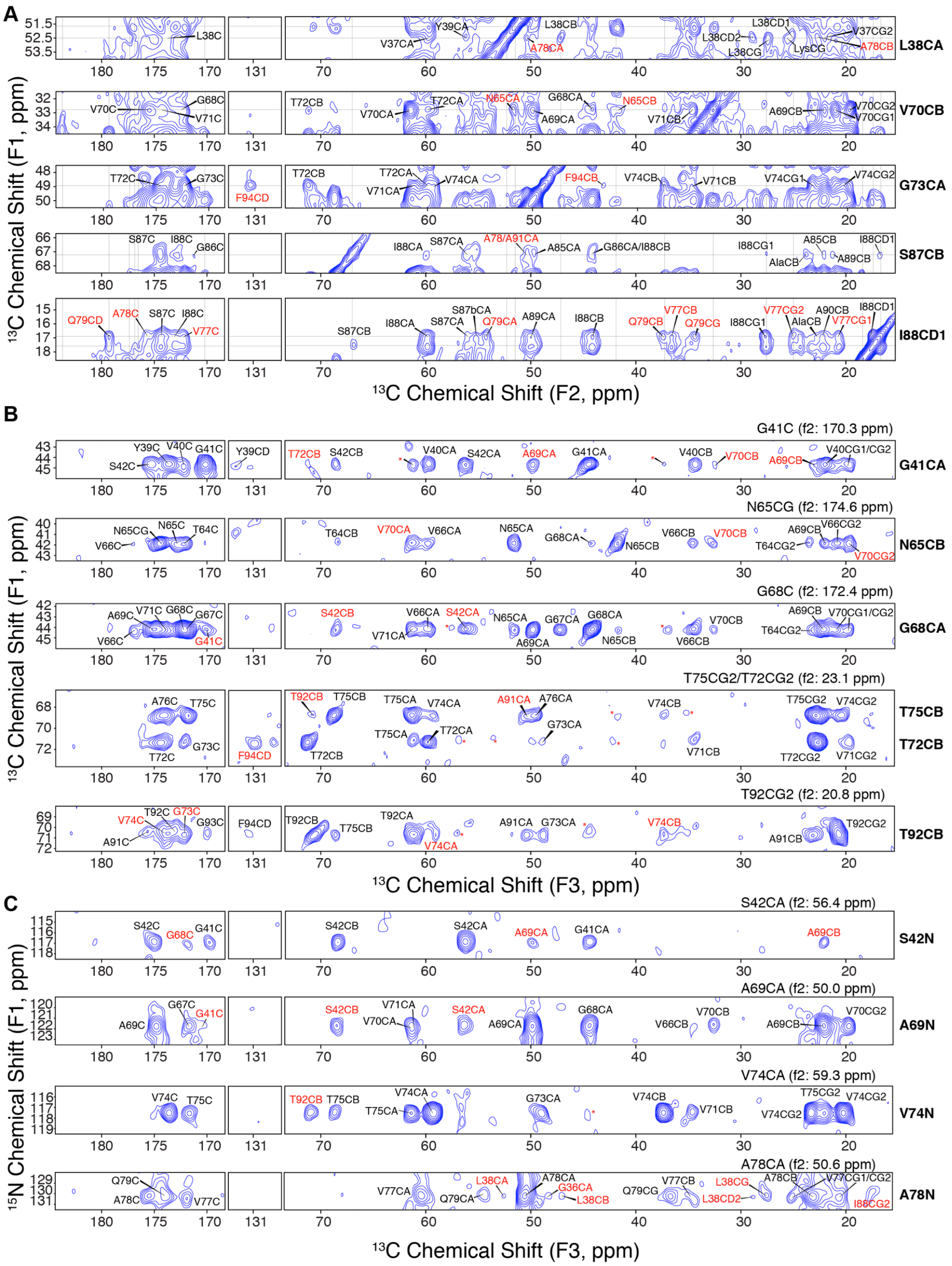

**Extended Data Fig. 6:** Long-range correlations detected in 2D and 3D PAR experiments. **a**, Strip plots with assignment labels from the spectrum in Extended Data Fig. 5 showing correlations from five atoms with unique assignments in F1. **b**, 2D strip plots from a 3D $^{13}C$-$^{13}C$-$^{13}C$ correlation spectrum showing correlations from six residues that are disambiguated in F1 and F2 relative to the 2D spectrum. **c**, 2D strip plots from a 3D $^{15}N$-$^{13}C\alpha$-$^{13}CX$ correlation spectrum showing correlations from four residues with unique $^{15}N$-$^{13}C\alpha$ shifts. All direct-dimension assignments are unique, arising from nearest neighbor residues or based on a network analysis. Long-range assignments are shown in red. Red asterisks indicate peaks that cannot be assigned based on the resonance list, cannot be disambiguated based on a network analysis, or are out of plane in the 3Ds.

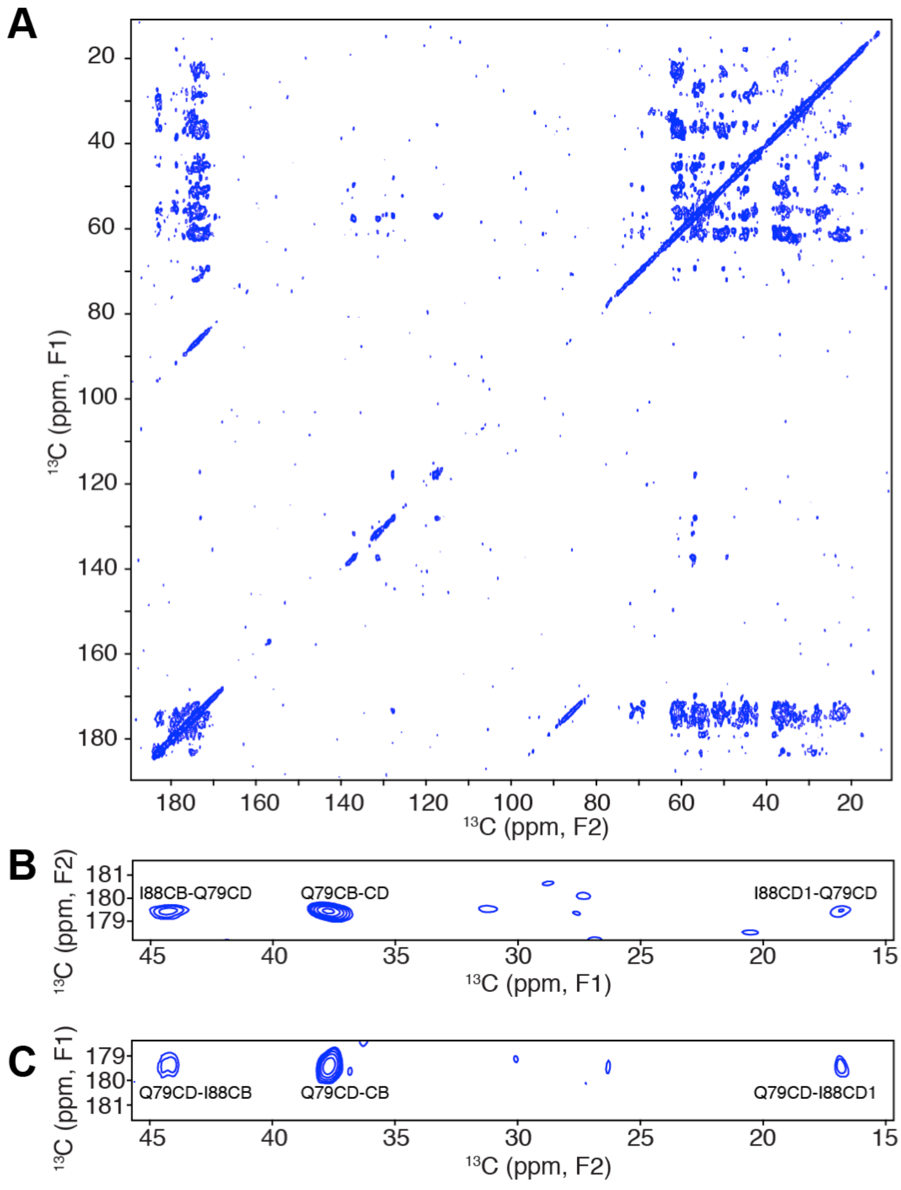

**Extended Data Fig. 7:** Two-dimensional $^{13}$C-$^{13}$C correlation spectrum acquired with DARR mixing on 25% [2-$^{13}$C]glycerol,$^{15}$N 75% natural abundance LBD Asyn fibril. **a**, $^{13}$C-$^{13}$C correlation spectrum showing the entire spectrum. **b**, Inset of I88-Q79CD correlations from upper left of diagonal. **c**, Inset of Q79CD-I88 correlations from lower right of diagonal. Data was acquired at 500 MHz $^{1}$H frequency with 11.111 kHz magic-angle spinning and a variable temperature set point of 0 ºC with 500 ms DARR mixing. The first contour is drawn at 5 times the root-mean-square noise.

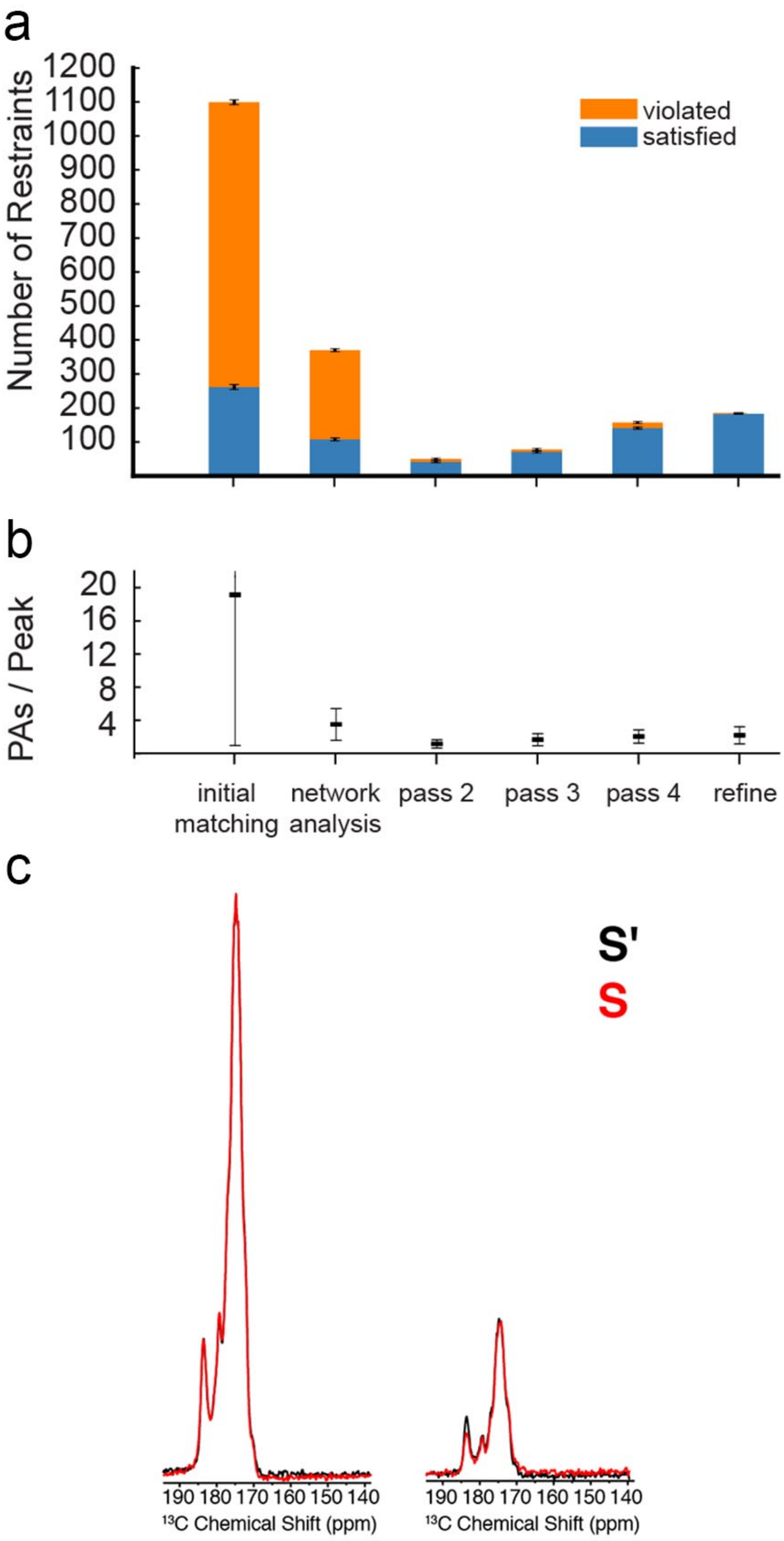
a
Number of Restraints
1200
1100
1000
900
800
700
600
500
400
300
200
100
violated
satisfied
b
PAs / Peak
20
16
12
8
4
initial matching
network analysis
pass 2
pass 3
pass 4
refine
c
S'
S
190 180 170 160 150 140
13C Chemical Shift (ppm)
190 180 170 160 150 140
13C Chemical Shift (ppm)

**Extended Data Fig. 8: PASD Results and FS-REDOR analysis. a**, Stacked bar chart showing the average number of restraints that are satisfied (blue) and violated (orange) by the top five lowest energy structures generated by the final refinement at each stage of the PASD structure calculation and refinement protocol. **b**, Average number of peak assignments (PAs) per peak at each stage of PASD. A high-likelihood cutoff of 0.90 was used. Error bars represent ± 1 standard deviation (n=11). **c**, Detection of salt bridges by FS-REDOR. $^{15}$N-dephased, $^{13}$C-detected frequency-selective REDOR (FS-REDOR) performed on uniformly $^{13}$C,$^{15}$N-labeled LBD Asyn fibril. Gaussian pi pulses were used to selectively reintroduce the dipolar couplings between the Lys NZ and the Glu CD resonances. The reference spectrum (S') is plotted in black and the REDOR dephased spectrum is in red (S) for short mixing (1.8 ms) and moderate mixing (9.0 ms).

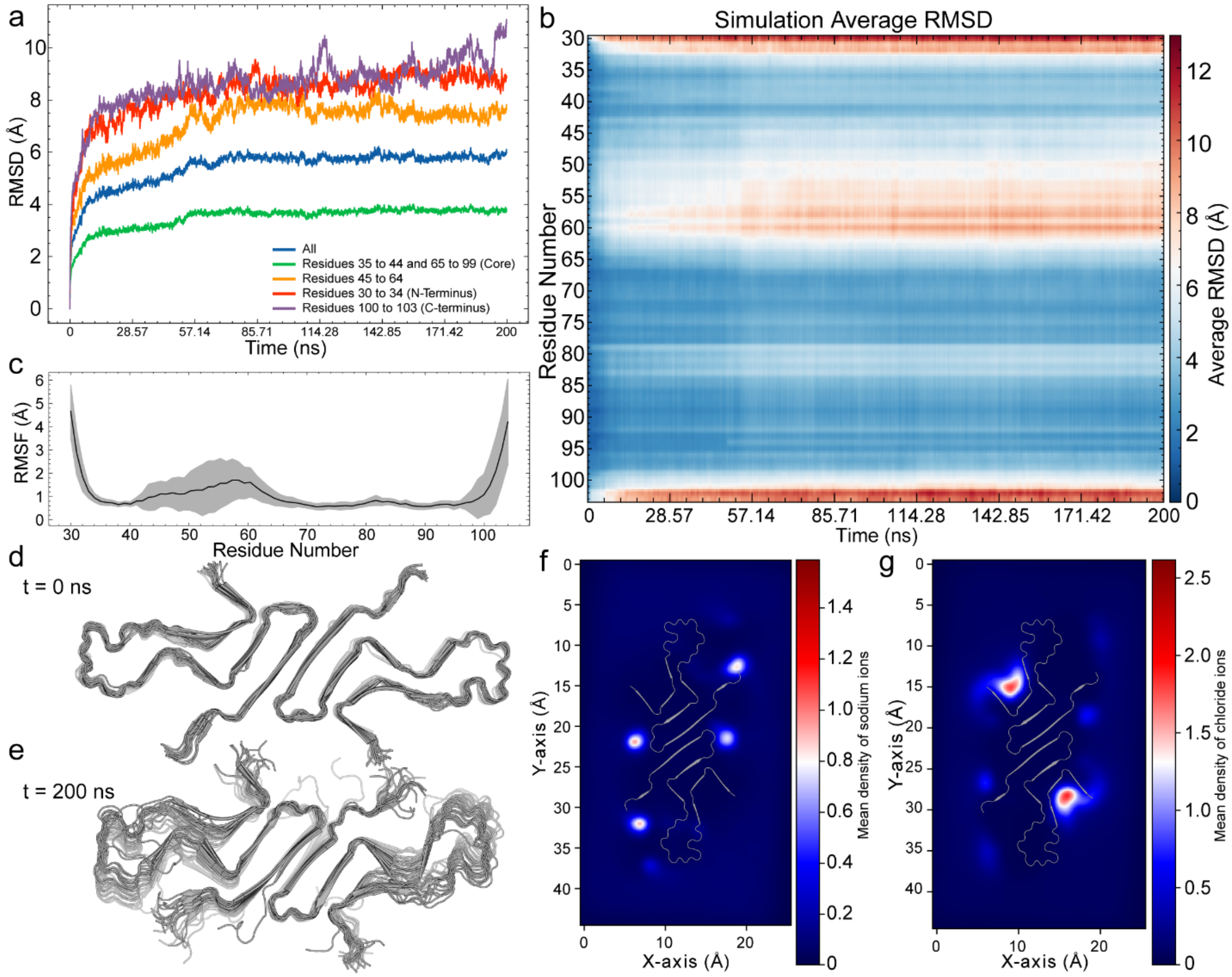

**Extended Data Fig. 9:** Unrestrained MD simulation (200 ns) of LBD fibril (2x 20-mer) from residues 30 to 103. Protomers corresponding to the top two and the bottom two protofilaments were removed as they do not replicate the extended nature of the fibril. **a**, RMSD of the full simulation time shown for all residues and specific portions of the structure. **b**, Residue specific RMSD averaged over each residue, n= 36. **c**, Residue specific RMSF of the fibril of the last 193 ns of the simulation, corresponding to where the RMSD in A and B plateau. **d-e**, Initial and final frames, respectively, of the simulation looking down the long axis of the fibril. **f**, Mean density distribution of sodium ions within the simulation, with four major sites arranged symmetrically across the protomers, coordinating to E83 and the C-terminus. **g**, Mean density distribution of chloride ions with a major coordination site between the lysines at residue 32, 34, 43, and 45, which are all pointing in towards the chloride ion density. This suggests that ions act as stabilizing forces regions of the fibril where charged residues stacked along the fibril axis and are not involved in salt bridge formation, neutralizing the charges present along the surface exposed residues along the long axis of the fibril.

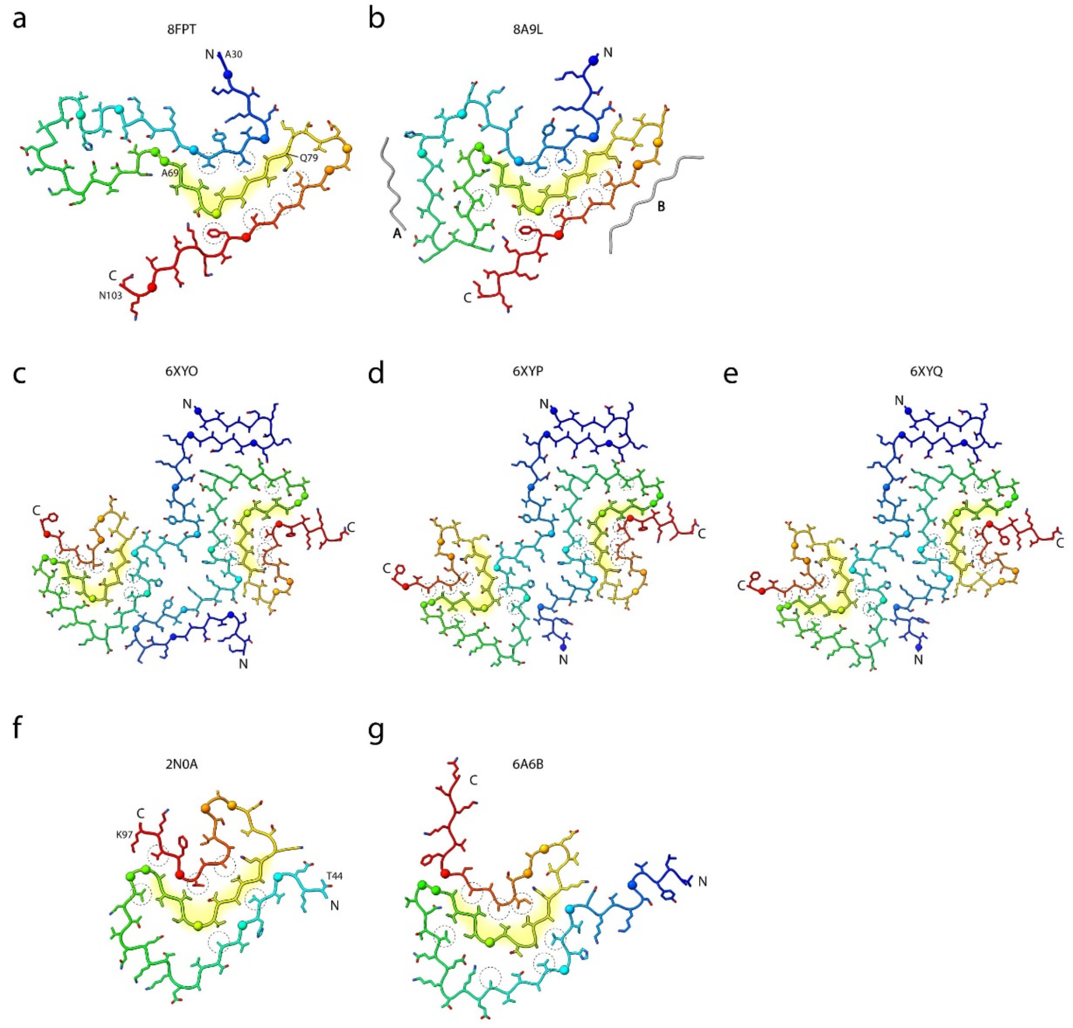

**Extended Data Fig. 10:** The conserved L-shaped motif in Asyn models. The L-shaped motif is comprised of residues 69-79 (highlighted in yellow) in the hydrophobic core of LBD amplified (**a**) and LBD extracted (**b**), MSA (**c-e**), and in vitro (**f-g**) Asyn fibrils. All models are colored by Jones' rainbow from N-terminus (blue) to C-terminus (red). Hydrophobic residues that interact with the L-motif are circled by a dotted line. The α-carbons of all glycine residues are depicted as a large sphere, and the unidentified proteinaceous islands 'A' and 'B' in the cryo-EM Asyn model (**b**) are in grey. Notably, the side chains pointing inside of the L-motif (A69, V71, etc.) interact with N-terminal strands (residues 35-42) in LBD structures, whereas they interact with C-terminal strands in MSA and in vitro structures (residues 86-94). Inversely, the side chains facing out of the L-motif interact with C-terminal strands (residues 86-97) in the LBD structures and with N-terminal strands in the MSA and in vitro structures (residues 41-66). The ability of Asyn to retain this embedded L-shape motif amongst such heterogeneity in folds is likely due to the high number of glycine residues.
